# Age-related Delays in Osteochondral Remodeling of Fracture Healing Illustrated by Mass Spectrometry Imaging

**DOI:** 10.64898/2026.02.13.705828

**Authors:** Charles A. Schurman, William Chandler, Diane Hu, Harrison Taylor, Nannan Tao, Theodore Miclau, Peggi Angel, Ralph Marcucio, Birgit Schilling

**Author notes:** Co-Corresponding Authors: Birgit Schilling, Ph.D. Professor, Buck Institute for Research on Aging Director, Mass Spectrometry and Proteomics Core Buck Institute for Research on Aging, Novato, CA 94945, USA, Ralph Marcucio, Ph.D. Professor Orthopaedic Surgery, School of Medicine, Department of Orthopaedic Surgery University of California San Francisco, San Francisco, CA 94110, USA.

## Abstract

Age-related delays in fracture healing are prevalent and contribute to morbidity and mortality in elderly populations. Clinical and preclinical studies demonstrate that aging is associated with slower and less complete fracture repair characterized by delayed cartilage and bone formation, impaired matrix resorption, and an increased risk of delayed union or non-union. Matrix-Assisted Laser Desorption/Ionization Mass Spectrometry Imaging (MALDI MSI) enables spatially resolved, in situ molecular analysis of proteins directly from murine fracture tissues. We applied collagenase type III (MMP-13) mediated proteolytic digestion to formalin-fixed, paraffin-embedded (FFPE) tibia fracture callus sections harvested 10 days post–tibial fracture from young (3-month-old) and aged (18-month-old) mice to perform spatially resolved proteomic profiling. Spatial MS Imaging revealed pronounced age-dependent differences in extracellular matrix protein composition and remodeling within the fracture callus. We identified upregulation of canonical bone and matrix proteins, including Col1a1 and Col1a2 specifically in the young fracture callus demonstrating advancement into harden callus formation. Conversely, Col2a1 and other soft callus proteins were only seen in the aged callus tissues. Further, protein indicators of tissue state, such as fibronectin (upregulated) and calreticulin (downregulated) were selectively regulated aged tissues, demonstrating a failure for aged tissues to fully progress into harden calluses. Spatial proteomic patterns demonstrated a marked delay in progression from cartilaginous to osseous callus in aged mice, consistent with impaired matrix remodeling during fracture repair. Together, these findings establish spatial MS Imaging based proteomics as a powerful approach to elucidate age-related alterations in fracture healing and to identify molecular regulators of impaired skeletal regeneration.

**Lay Summary:** Using spatially-resolved proteomics via mass spectrometry imaging on fracture callus tissues from young and aged mice, we observed delayed healing in aged animals based on the composition of the extracellular matrices. Higher levels of bone specific collagens were detected in young animals, whereas cartilage specific collagens were detected in aged animals at higher levels. Further, detection of novel, non-canonical callus proteins revealed critical transitional steps that are delayed in aged-callus tissues, and these may also contribute to the delayed healing aged animals.

## Introduction

Age-related delays in bone fracture healing in the elderly are common, and these fractures are associated with a high rate of morbidity and mortality^1,2^. In most healthy adults, fracture healing progresses normally over the course of about 6-8 weeks and usually does not significantly impact quality of life^3^. However, in elderly patients the time of recovery after fracture is a significant cause of morbidity and mortality^4,5^. Bone fractures heal through two distinct processes. In fractures that are rigidly stabilized, bone forms directly through intramembranous ossification. In contrast, fractures that are not stabilized stem cells differentiate into chondrocytes that make a large cartilage callus that transforms into bone through the process of endochondral ossification^6^. Numerous studies document a slower and less complete healing of bone fractures in older individuals, compared to younger individuals^7–12^, and animal studies show that endochondral and intramembranous ossification are delayed in elderly animals^9,13,14^. Additionally, older fracture patients are at higher risk for delayed union or non-union of fractures that require additional surgical intervention. Even when healed, older individuals experience diminished bone volume and decreased mechanical strength with a 15–25% decrease in structural stiffness and force to refracture as shown in animal models, leading older individuals to be higher risk for refracture^7,15,16^. While the time for healing is individualized and depends on numerous comorbidities^17,18^, delayed fracture healing represents a large clinical challenge, and treatments that accelerate repair are an unmet clinical need especially for a world-wide aging population. Understanding the molecular regulators of the healing process will serve to develop novel therapeutic interventions and lower morbidity and mortality in the elderly.

Several mechanisms that partially contribute to delayed fracture healing in the elderly have been identified. First, aging leads to chronic, elevated inflammation (“inflammaging”)^19^ and disturbed immune responses, which in turn impair cellular recruitment to the callus, proper osteogenic differentiation, and the production of cytokines necessary for healing^7,20–26^. Rejuvenating the immune system stimulates healing in aged mice^27,28^. How this works is not known, but the extended presence of inflammatory factors such as Interleukin 22 (IL22) can prolong the inflammatory response that precedes callus formation. Prolonged inflammation can impact the expression of genes that are critical modulators of healing including BMP-2, Indian Hedgehog (IHH), VEGF, and COX-2, in aged fracture calluses which can slow chondrocyte and osteoblast differentiation^8,9,16,29^ from already depleted osteochondral stem cells in older tissues^30^. Macrophages may also play a role in age-related changes in fracture healing^31^, and they may have changes in signaling systems that lead to pro-inflammatory states^32^. Further, aged tissues often exhibit increased senescent cell burden and oxidative damage which accumulates with age, and following injury, impairs local tissue regeneration and immune responses, further hindering repair processes^21–23^. Additionally, decreased blood flow and angiogenesis occur at fracture sites in older individuals, delaying the delivery of nutrients and signaling molecules essential for bone healing^10,16,33^. It is also thought that aging reduces extracellular matrix (ECM) protein production, such as collagens, and their post-translational modifications (cross-linking) that may weaken bone and lower or slow mineralization of the callus^10,34^, though less is known specifically about ECM regulation during fracture callus remodeling or how this is disrupted with age.

Matrix-Assisted Laser Desorption/Ionization Mass Spectrometry Imaging (MALDI MSI), introduced in the 1990s^35,36^, enables spatially resolved detection of analytes directly from tissue sections, providing an alternative to conventional histology by imaging proteins, lipids, metabolites, and glycans in situ^37–41^. Applied across diverse disease contexts, including cancer, neurodegeneration, and others^42–45^, spatial MS Imaging has recently begun to be leveraged to study musculoskeletal tissues, including bone, cartilage, and the synovium revealing roles for lipid species in inflammation and identifying region-specific glycan alterations linked to mechanical loading Osteoarthritis (OA) ^46–57^. While effective for small molecules and glycans, use of MS Imaging for proteomic applications to the skeleton have been limited to softer tissues such as cartilage and the synovium^58,59^. Advancements in spatial MALDI MS Imaging, utilizing proteolytic-enzymatic digestions, have allowed for the targeted analysis of matrix proteins^44,45,60,61^. Specifically, pre-treatment with collagenase type III (Col-III / MMP-13) enables spatial-proteomic analysis of the dense, crosslinked extracellular matrix of bone and cartilage^62^. This enzyme targets collagens I–III and regulates ECM turnover. Recently, we demonstrated that collagenase III / MMP13 pretreatment provided excellent molecular detail of subchondral bone remodeling in human osteoarthritic knees^62^. Given that the healing fracture callus is a mosaic of dynamically regulated ECM materials, primarily collagenous cartilage and bone, Col-III–assisted spatial proteomics via spatial MS Imaging offers an exciting opportunity to reveal new molecular details of fracture callus remodeling.

The objective of this study was to identify specific changes in the ECM of the fracture callus undergoing endochondral ossification during healing in young and aged animals to identify and visualize barriers to healing in aged mice using sections from animals that we have previously shown to have delayed in healing^9^. Generating spatial maps of tissue remodeling in the fracture callus during healing, and through identification of specific matrix proteins, provides molecular evidence and visual confirmation of specific barriers in endochondral remodeling that may contribute to delayed healing in aged animals. Identification of specific collagens and new matrix-associated proteins within the callus of young and aged mice at important transitional regions in the callus provide potential new molecular regulators of the healing process which may be targets for future research. In conclusion, we demonstrate the first use of ECM-targeted spatial proteomics via spatial MS Imaging in the endochondral fracture callus and identify potential regulators of callus remodeling that are absent or delayed in aged tissues.

## Methods

### Animals

Male C57BL/6J mice were purchased from Jackson Laboratory and housed and bred at the University of California San Francisco. All animal procedures were conducted in accordance with the UCSF Institutional Animal Care and Use Committee (IACUC) guidelines and were approved under protocol number [IACUC #03024337-03D]. Mice were wild type, with no genetic modifications. Mice were aged to 12 weeks (3 months) and 18 months of age. Animals were maintained behind a barrier and were immunocompetent and healthy at the time of study. Mice were housed 2–5 mice per cage / single housing in individually ventilated cages under Specific Pathogen Free (SPF) conditions. Rooms were maintained on a 12:12 h light–dark cycle at a temperature of 20–26 °C and relative humidity of 40–60%. Standard rodent chow (LabDiet 5001, Purina, product #0001328) and autoclaved water were provided *ad libitum*. Bedding and nesting material were supplied for environmental enrichment, and cages were changed according to facility husbandry protocols.

Prior to fracture, mice were anesthetized with 2’X avertin. Tibial fracture was induced and post-treated in a manner designed to produce endochondral remodeling^9,63^. Briefly, closed fractures of the mid-tibia were created by three-point bending, fractures were not stabilized, and animals were allowed to move freely for 10 days before collection. Buprenex (0.1 mg/kg) was given after fracture and when animals were in pain. No unexpected severe adverse events were observed in any experimental group. Minor anticipated adverse effects included transient weight loss <10%, mild post-procedural discomfort, and temporary reduction in activity, which resolved without intervention. No mortality occurred outside of planned humane endpoints. To minimize adverse events, animals were monitored daily for health and behavior. If early signs of distress were observed (e.g., ruffled fur, reduced mobility, >15% weight loss), animals were evaluated by veterinary staff and removed from the study if humane endpoint criteria were met.

### Preparation of FFPE Mouse Bone Fracture Tissue Sections

The tibiae were dissected at day 10 post injury and the skin was removed. Tissue was fixed in fresh 4% PFA in PBS (pH 7.4) overnight (12–18 hours) at 4 °C with gentle rocking. After fixation, samples were rinsed 3 × 10 minutes in PBS at 4 °C followed by decalcification in 19% ethylenediaminetetraacetic acid (EDTA) for 14 days with gentle agitation at 4 °C, replacing the EDTA every three days until the bone appeared semi-transparent while maintaining internal structure. Specimens were then dehydrated through a graded ethanol series, cleared with xylene, and then embedded in paraffin lengthwise with the sagittal plane horizontal to the face of the block. Sections (5–7 µm) were cut using a microtome (Leica Microsystems) and mounted on glass microscopy slides.

### Preparation of FFPE Fracture Callus Tissue for Spatial Proteomics

FFPE fracture callus sections were processed for enzymatic digestion using adapted protocols to accommodate the extensive extracellular matrix^40,64–66^. Slides were incubated at 60 °C for 1 h to soften paraffin, then deparaffinized with two 3 min xylene washes. Sequential rehydration included two 1-minute washes in 100% ethanol, one 1-minute wash in Carnoy’s solution (60% ethanol, 30% chloroform, 10% glacial acetic acid), 1-minute washes in 95% and 70% ethanol, and two 3-minute washes in distilled water. Antigen retrieval was performed in citraconic anhydride buffer (pH 3.0; Thermo) using sealed slide mailers incubated overnight at 60 °C (≤ 12 h) while fully submerged in an external buffer reservoir. Buffers were exchanged gradually with water to avoid tissue disruption after cooling to RT. Deglycosylation was achieved by applying PNGase F (0.1 µg/µL; N-Zyme Scientifics) with a TM Sprayer M3 (HTX Imaging), followed by a 2 h incubation at 37 °C in humid chambers. Released N-glycans were washed away with a series of washes including 70% ethanol, followed by alternating washes in high-pH (10 mM Tris, pH 9) and low-pH (10 mM citraconic anhydride, pH 3) buffers, ending with distilled water. For ECM-targeted, collagenase digestion, a second antigen retrieval was performed (10 mM Tris, pH 9, overnight at 60 °C). Collagenase III (0.1 µg/µL; Worthington Biochemical) was applied via TM Sprayer M3 and incubated for 5 h at 37 °C in humid chambers. After drying, slides were coated with CHCA matrix containing [Glu1]-fibrinopeptide B (200 fmol/L; Sigma-Aldrich), dried for 30 min in a vacuumed desiccator, briefly immersed (< 1 s) in cold 5 mM ammonium phosphate monobasic, and immediately desiccated for 1 min (at most).

### Spatial Mass Spectrometry Imaging

Matrix-assisted laser desorption/ionization mass spectrometry imaging (MALDI MSI) was performed using a trapped ion mobility spectrometry–quadrupole time-of-flight instrument (timsTOF fleX; Bruker) in positive ion mode. Acquisition settings were: m/z range 600–2500, plate offset 70 V, deflection 60 V, funnel 1 RF 400 Vpp, funnel 2 RF 500 Vpp, multipole RF 400 Vpp, transfer time 78 µs, pre-pulse storage 20 µs, collision cell energy 25 eV, collision RF 2500 Vpp, quadrupole ion energy 10.0 eV. Calibration was performed before each run (Agilent G1969-85000 standard). Imaging was conducted with a 20 µm pixel resolution, 300 laser shots per pixel, and a 14 µm x 14 µm laser application (18 µm x 18 µm as resulting field size).

### Hematoxylin and Eosin Histological Staining

After imaging, the matrix was removed using 100% ethanol, followed by H&E staining using Gill‘s 2 hematoxylin 2 and Eosin-Y Fisher. Tissues were rinsed for 30 seconds in 95% ethanol, then 70% ethanol for 30 seconds each, followed by a 2 minute incubation in Hematoxylin to stain nuclei. Excess stain was removed through two water washes at 2 minutes each, followed by 95% ethanol and 100% ethanol for 30 seconds each. Eosin-Y was used to stain cytoplasm and extracellular composition for 30 seconds and excess was removed through 3 rinses in 100% ethanol. Slides were dried any coverslips were mounted using Cytoseal. Images were digitally recorded using a Hamamatsu NanoZoomer.

### Data Processing and Statistical Analysis

Mass spectrometry data were processed in SCiLS Lab MVS v2025a Pro (Bruker Scientific). Spectra were background-normalized and resampled to a common mass axis using a total-ion-count-preserving algorithm. All spectra were further normalized to the [Glu1]-fibrinopeptide B standard (Sigma F3261) at m/z 1570.6768. The top 500 most intense m/z features per sample were retained to reduce noise while maintaining biological variability and used for downstream processing. Initial group-wide analysis was completed by defining age groups as classes: 3 month-old animals (Young) vs 18-month-old animals (Aged). Later, spatial segmentation was performed using bisecting k-means clustering utilizing Manhattan distance to identify spatial regions with similar molecular identity. The largest spatial regions of interest (ROIs) were generated using hierarchical segmentation. The two largest classes, Green (“Bone-like”) and Yellow (“Cartilage-like”) ROIs were selected for use in Discriminating Feature analysis in the SCiLS lab. In both “Young vs Aged” and “Bone-like vs Cartilage-like” analyses, candidate features were identified using the receiver operating curve (ROC), with inclusion criteria of area under the receiver operating curve (AUROC) ≥ 0.7 as the statistical threshold for significant class prediction. For identification of candidates in the “reverse” direction (e.g. upregulated in aged vs young) an AUROC cutoff of |AUROC| < (1 - 0.7) = 0.3 was used for significance. Candidate m/z values were validated across the eight independent specimens by exporting ROI-specific intensity data and calculating mean intensity for each independent ROI for statistical comparisons. For analysis using the automatically generated tissue regions (Bone-like vs Cartilage-like) sub-regions from each sample were created producing regions with dual annotations of Young “Bone-like”, Young “Cartilage-like”, Aged “Bone-like”, or Aged “Cartilage-like”. Feature matching for candidate features was completed using a collagenase-digested mouse bone library^67^. Identifications were accepted with a mass tolerance ≤ 2 mDa (∼ +/- 15 ppm). Statistical comparisons of ROI mean intensity were completed in GraphPad Prism v10.1.1 with an *a priori* set alpha of 0.05 (p < 0.05) for all comparisons. For the Young vs. Aged analysis, a one-sided Student’s t-test was used. For analysis of feature intensity for “Bone-like” vs. “Cartilage-like”, a paired one-sided Student’s t-test was used with sample ID as the linking factor for paired ROIs. A one-sided t-test was selected, given that the direction of regulation for a given comparison was known, as determined by the previous AUROC statistical test. For statistical comparison for the reclassified regions following the tissue gradient (Figure 6), a one-way ANOVA with Bonferroni corrected *post-hoc* pairwise comparisons was used with factors as the new “gradient” tissue regions (I, II, III).

### Data Availability

The spatial proteomic datasets have been uploaded to the Center for Computational Mass Spectrometry MassIVE repository at UCSD and can be accessed using the following link: ftp://massive-ftp.ucsd.edu/v11/MSV000100309/ (MassIVE ID number: **MSV000100309**)

(https://massive.ucsd.edu/ProteoSAFe/dataset.jsp?task=6a4584b068564b7990287c895a0f148a)

## Results

Age-related delays in fracture healing result from multiple molecular complications, including alterations in the production of extracellular matrix (ECM) proteins^10,34^. To visualize ECM proteins during fracture healing, Matrix Assisted Laser-Desorption Ionization Mass Spectrometry Imaging (MALDI MSI) was used to analyze the ECM composition of murine formalin fixed paraffin embedded (FFPE) fracture callus tissue sections. This application of spatial MS Imaging is the first to employ spatial proteomics based on an enzymatic digestion of endochondral tissue from mouse fracture callus. This process involves the application of an ECM-specific collagenase III to deparaffinized and rehydrated FFPE tissues followed by MALDI matrix (α-cyano-4-hydroxycinnamic acid, CHCA) through controlled spraying directy onto tissue slides (**Figure 1A**). Prepared slides are then imaged with the timsTOF fleX mass spectrometer (**Figure 1B**), which collects single pixel mass spectra as discrete locations across the tissue sections in a raster manner. These single pixel mass spectra are then reassembled and extracted to build spatial heat maps corresponding to the intensity of molecular features (proteins and corresponding peptides) across the imaged tissue while imaged tissue sections are recollected and utilized to generate tissue-specific peptide libraries for feature identification (**Supplemental Figure 1**).

**Figure 1:**
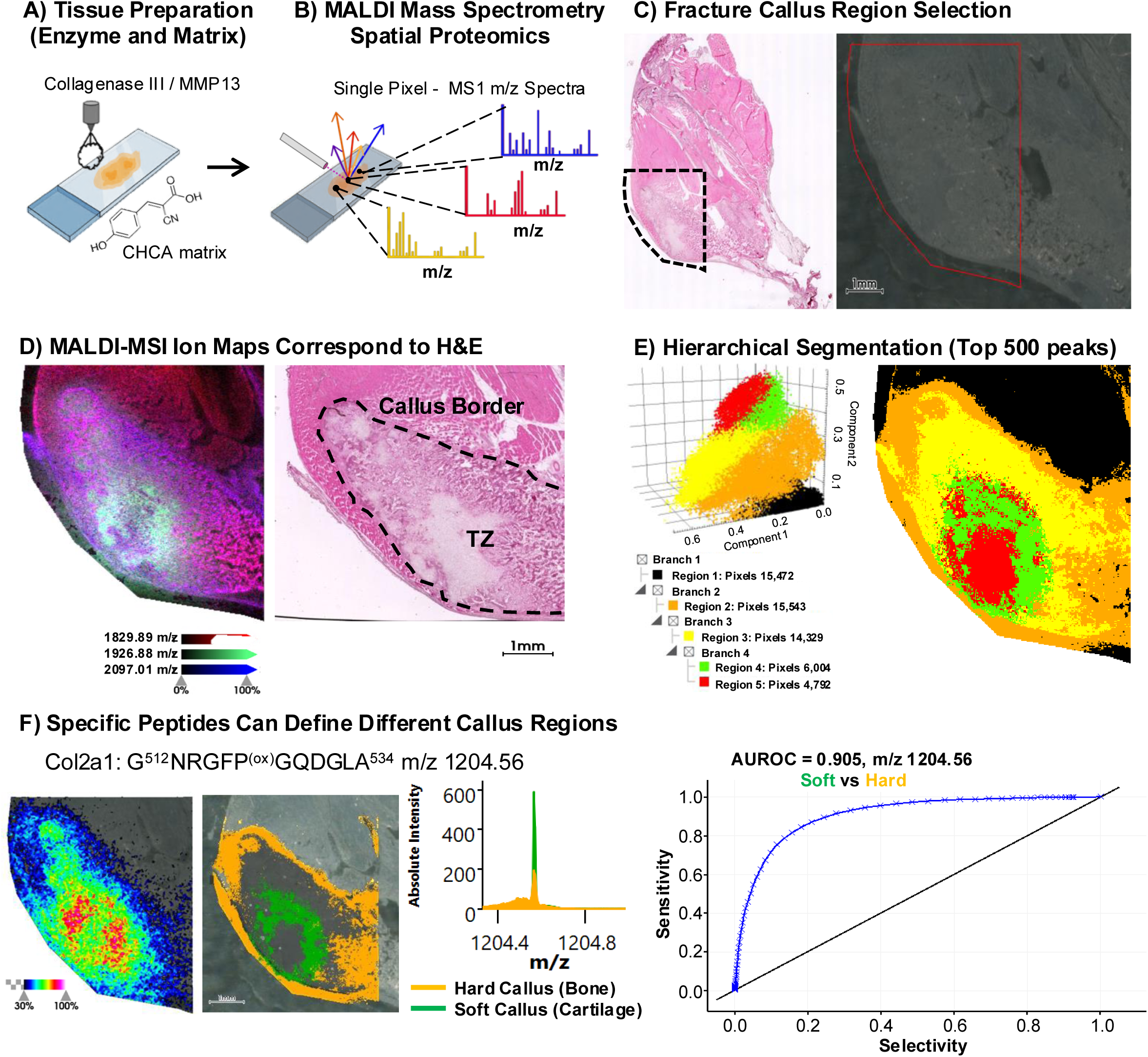
MALDI MS Imaging interrogates Spatial Differences in the Fracture Callus. **A)** FFPE tissue slides of fracture callus were prepared for Matrix Assisted Laser Desorption/Ionization (MALDI) Mass Spectrometry Imaging (MSI) via serial enzymatic digestion with PNGase F and Collagenase III followed by application of α-Cyano-4-hydroxycinnamic acid (CHCA) matrix via the HTX-M5 sprayer. **B)** Single pixel mass spectra were collected with the Bruker timsTOF fleX across **C)** segmented callus regions as identified through hematoxylin and eosin (H&E) staining of serial tissue sections. **D)** H&E staining of the callus shows distinct tissue types in the fracture callus including the callus border and the transition zone (TZ) between cartilaginous and boney regions. These histologic features are well captured by molecular signal of ion features **(E)** detected via MALDI MSI. Hierarchical segmentation **(F)** based on the top 500 most abundant peaks in the MALDI spectra reveal distinct separation of regions of tissue within the fracture callus that align to morphologic features via H&E. Quantification of a known peptide of the cartilage marker protein Col2a1 **(G)**, G^513^NRGFP^(ox)^GQDGLA^524^ at m/z = 1204.56, showed increased signal in the green segment around the soft cartilaginous callus (green) compared to the outer hard callus region (orange) such that the area under the receiver operating curve (AUROC) = 0.905.

Young 3-month-old (N=4) and aged 18-month-old (N=4) C57Black/6 mice received tibial fracture and were left to heal without fixation (mechanically unsupported) for 10 days. After 10 days, hind limbs were collected and prepared for FFPE processing and tissue sectioning. To capture the complex mixture of tissue types including bone, cartilage, muscle, and other soft tissues surrounding the fracture callus, entire unstained tissue sections of the hind limb were imaged with spatial MS Imaging after collagenase digestion. After imaging, slides were collected and stained with Hematoxylin and Eosin (H&E) and the region of tissue containing the healing callus was targeted for down-stream analysis of spatial MS Imaging spectral data (**Figure 1C, Supplemental Figure 2**). Selected features (peptide ions) from MS Imaging could accurately recapitulate physiologic features found in H&E staining of the tissues, including identification of the callus border and the transition zone (TZ) between soft and hard callus regions (**Figure 1D**). This demonstrated that spatial MALDI MS Imaging accurately captured the molecular complexity and gradient of tissue remodeling in the healing fracture callus. Further, unbiased quantitative tools, such as a Hierarchical Segmentation (**Figure 1E**) acting on hundreds of molecular features, were able to identify distinct tissue regions of interest (ROIs) that had a shared molecular identity and matched histological features. These automatically defined ROIs were used for the quantification of the abundance of individual features within and between different tissue segments allowing for precise knowledge of the location and abundance of different peptides derived from ECM proteins across the callus. For example, the ion heatmap or spatial distribution for a known peptide of Collagen Type II, G^512^NRGFP^(ox)^GQDGLA^524^ m/z = 1204.56 is shown and its abundance quantified within the automatically generated Green and Orange regions of interest (ROIs) that spatially correspond to the soft, cartilaginous and hard, bony callus regions, respectively (**Figure 1F**). Receiver Operating Curve (ROC) analysis can also be used to determine if the spatial distribution of specific features was statistically predictive for one ROI compared to another. For instance, the shown peptide for Col2a1, had a statistically significant (>0.7) area under the receiver operating curve (AUROC) value of 0.905 when comparing its distribution in the green (soft callus) ROI against the orange (hard callus) region.

In order to first assess the differences in the fracture calluses in young vs aged animals (**Figure 2A**), spectral data for all eight animals (young 3-month-old (N=4) and aged 18-month-old (N=4) mice) were analyzed simultaneously as one composite data set. Immediately, it became evident that the MS Imaging spectra (**Figure 2B**) corresponding to the young and aged samples exhibited a rich molecular landscape that differed from each other in the abundance of numerous peaks, each of which corresponded to separate molecular features/peptides. Using the categories of “young” and “aged”, a discriminating feature analysis utilized the receiver operating curve (ROC) and was utilized to detect mass spectrometric peaks or features that best described the difference between the young and old fracture callus tissue samples. Using an area under the receiver operating curve (AUROC) cutoff of >0.7, 31 unique m/z features were found that were either upregulated and predictive for tissue samples from the young (with 27x m/z features) or aged (with 4x m/z features) mice (**Supplemental Table 1**). Hierarchical clustering of each individual sample by m/z feature intensity revealed that tissue sections from young and aged mice clustered separately reflecting distinct molecular differences (**Figure 2C).** Matching of these m/z features to a collagenase digested spectral library^67^ of mouse bone and cartilage tissues identified several m/z features as peptides from known skeletal ECM proteins, namely Collagens 1, 2, and 6 as well as 2 separate peptides derived from Calreticulin (Calr). The uniqueness of molecular signatures between age groups was further exemplified in a Principal Component Analysis (PCA) of these features, distinguishing fracture calluses from young and aged mice on the molecular level (**Figure 2D**). The abundance of each individual feature was also further statistically tested on the cohort level using a t-test (**Supplemental Figure 3**), and 29 of the 31 features showed significant differences in abundance between calluses from the young and aged mice. Of the 31 identified features, the features with the greatest difference included two features upregulated in the tissues from young animals - m/z 1829.90 and m/z 1637.79 - and two m/z features upregulated in tissues from the aged animals - m/z 1435.79 and 596.31 (**Figure 2E**). Strikingly, feature m/z 1435.79 which was upregulated in callus tissue from aged mice, matched a peptide of Col2a1 G^1101^PQGLAGQRGIVGLP^ox-1112^, was the only identified peptide of Col2a1 in this comparison. On the other hand, the feature m/z 1637.79 upregulated in the callus from young mice matched to a known peptide of Col1a1 G^1044^PVGPAGKN^de^GDRGETGPA^1061^. In fact, all identified peptides derived from collagen type 1 were only upregulated in the callus tissues from young mice after 10 days of healing. In our analysis, the only other ECM protein identified and quantified as significantly different in our analysis between the age groups was Calreticulin (Calr), a calcium chelator protein that can also act as an osteoblast proliferation factor. Calr was identified through two unique peptides (**Supplementary Figure 4**) and was detected only in the callus tissues from young mice.

**Figure 2:**
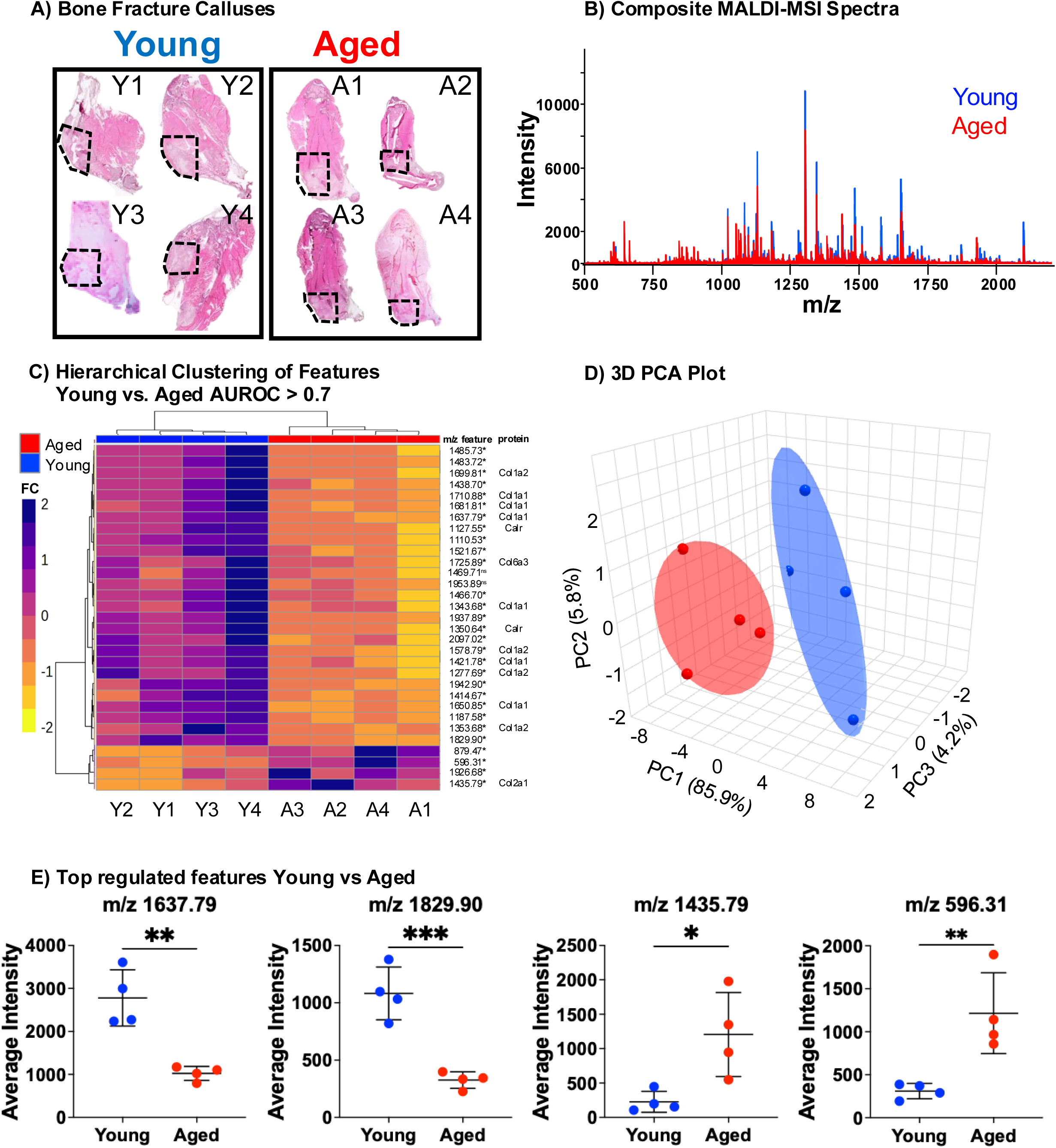
Molecular Differences in Young and Aged Callus Detected by MALDI MSI. **A)** Fracture callus from Young and Aged (N=4 ea.) mice 10 weeks post fracture were collected and prepared for FFPE sectioning, H&E staining, and callus subregions were selected for MALDI MSI. **B)** The mean mass spectra for the young (blue) and aged (red) groups are overlain, showing rich and diverse signal for many molecular features that may distinguish young and aged tissue. Discriminating Features analysis in SCiLS identified 31 unique m/z features that described the difference between the combined young and aged tissues with an |AUROC| > 0.7. Hierarchical clustering of intensity for each of these 31 features from each tissue region showed young and aged samples cluster separately **(C)** and PCA analysis **(D)** of these features similarly showed group separation between young and aged samples. **(E)** Scatter plots for two features up-regulated in young tissue, 1829.90 m/z and 1637.79 m/z, and two features up-regulated in aged tissue, 1435.79 and 596.31 m/z, are shown that display significant difference when feature intensities from each tissue section at that m/z are compared with student’s t-test. *p<0.05, **p<0.01 ***p<0.001

Visualizations of the selected top-regulated features are shown in **Figure 3**. Two peptides derived from Col1a1 with m/z 1829.89 (**Figure 3A**) and 1637.79 (**Figure 3B**) showed significantly increased intensity in the callus sections from young mice (top rows). The AUROC score for both peptides was above the statistical threshold for prediction (>0.7) for the “young” tissue regions compared to the “aged” group (bottom rows), with AUROC values of 0.902 and 0.936 respectively. Features m/z 1437.79 (**Figure 3C**) – previously identified as Col2a1 – and an unidentified, novel feature m/z 596.31 (**Figure 3D**) demonstrated significant prediction for the “aged” region with AUROC values of 0.201 & 0.078 respectively. This indicated that the callus tissues from the aged animals did contain some unique, predictive identifier peptides that were absent from the callus sections of the young mice.

**Figure 3:**
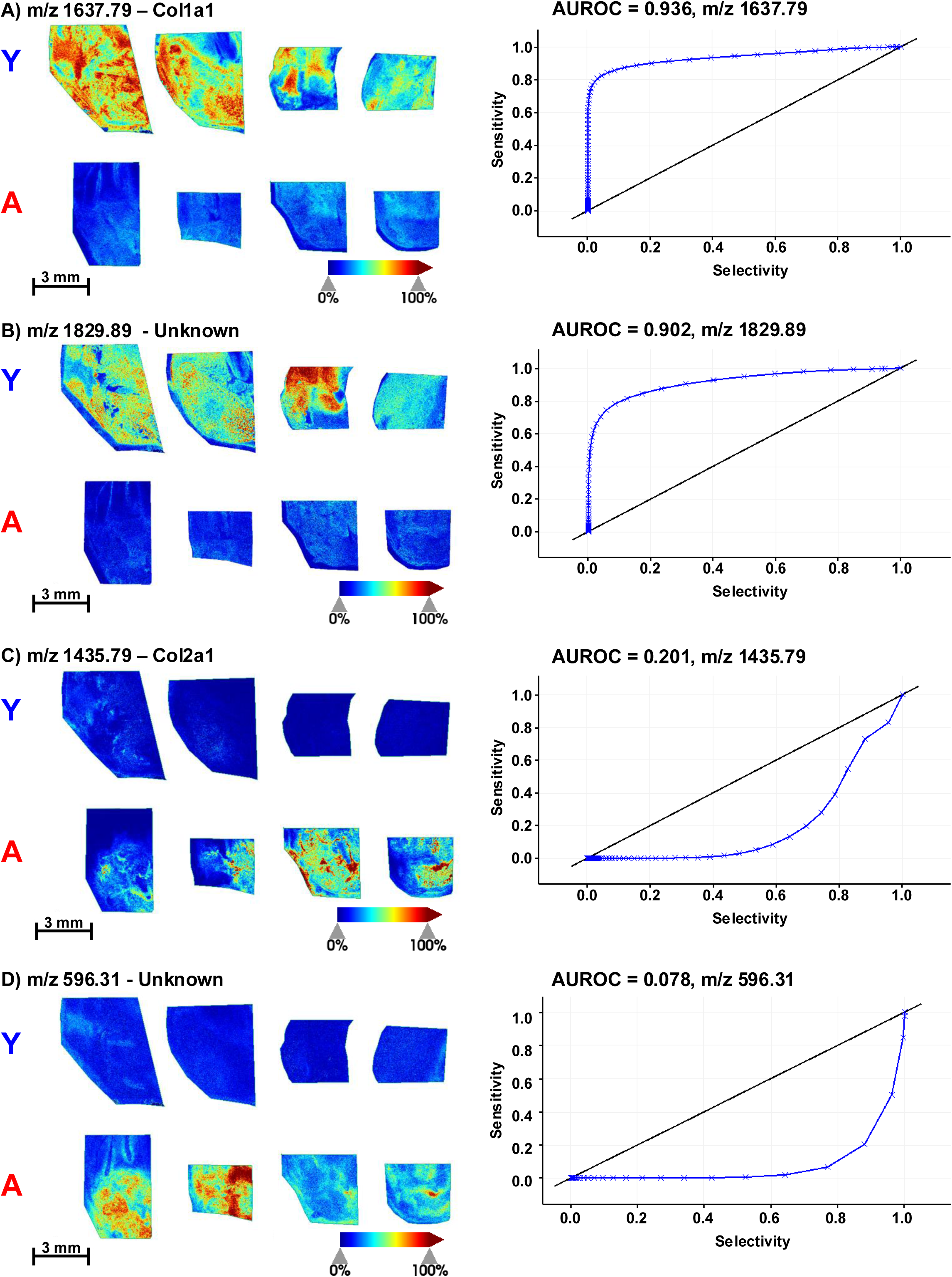
Spatial Heat Maps for Regulated Features in Young and Old Fracture Callus. Spatial Heatmaps for molecular features with significant regulation by age group in healing fracture calluses. Two features, m/z 1829.89 **(A)** and m/z 1637.79 **(B)** showed higher intensity in young fracture calluses (top) with very little or no noticeable signal in aged samples (bottom). These two features have an AUC >0.9 in discriminating feature analysis utilizing the receiver operating curve indicating that the presence of these features had highly significant predictive ability for the fracture callus from young animals compared to the aged fracture callus tissue. Conversely, two other selected features had inverse regulation and were upregulated in the aged fracture callus compared to young. These features, m/z 596.31 **(C)** and m/z 1435.79 **(D)**, had AUC values less than 0.3 in discriminating feature analysis utilizing the receiver operating curve, the statistical cutoff for significant prediction for the aged fracture callus tissue.

Collagen Type I and Collagen Type II are often used to delineate bone from cartilage and two of the top regulated peptides in between the callus sections from young and aged mice were derived from these proteins. Thus, we sought to determine in the same tissue sections if these proteins could be spatially mapped to identify the progression of callus healing across the age groups. First, additional peptides for Col1a1 (m/z 1483.72 G^450^PAGEEGKRGARGEP^(ox)-464^) and Col2a1 (m/z 1907.87 G^513^NRGFP^(ox)^GQDGLA^524^) were identified from our tissue-specific spectral library. Then, in a representative fracture callus from a young mouse, spatial maps were generated for each peptide independently (**Figure 4A**). Intensity for the Col1a1 peptide (indicator for bone tissues) was strongest at the outer edges of the callus where the furthest progression of tissue healing towards bone matrix would have occurred. Col2a1 (indicator for cartilage tissues) showed the strongest signal towards the center of the callus where the soft, cartilaginous callus would take the longest to remodel. Overlay (**Figure 4B**) of the signal from each peptide more clearly delineated the boundary between the different matrix regions and illustrated the transition between tissue types as the callus turned over from cartilage (pink) to bone (blue) with almost discrete separation in areas where the two peptides had the highest abundance.

**Figure 4:**
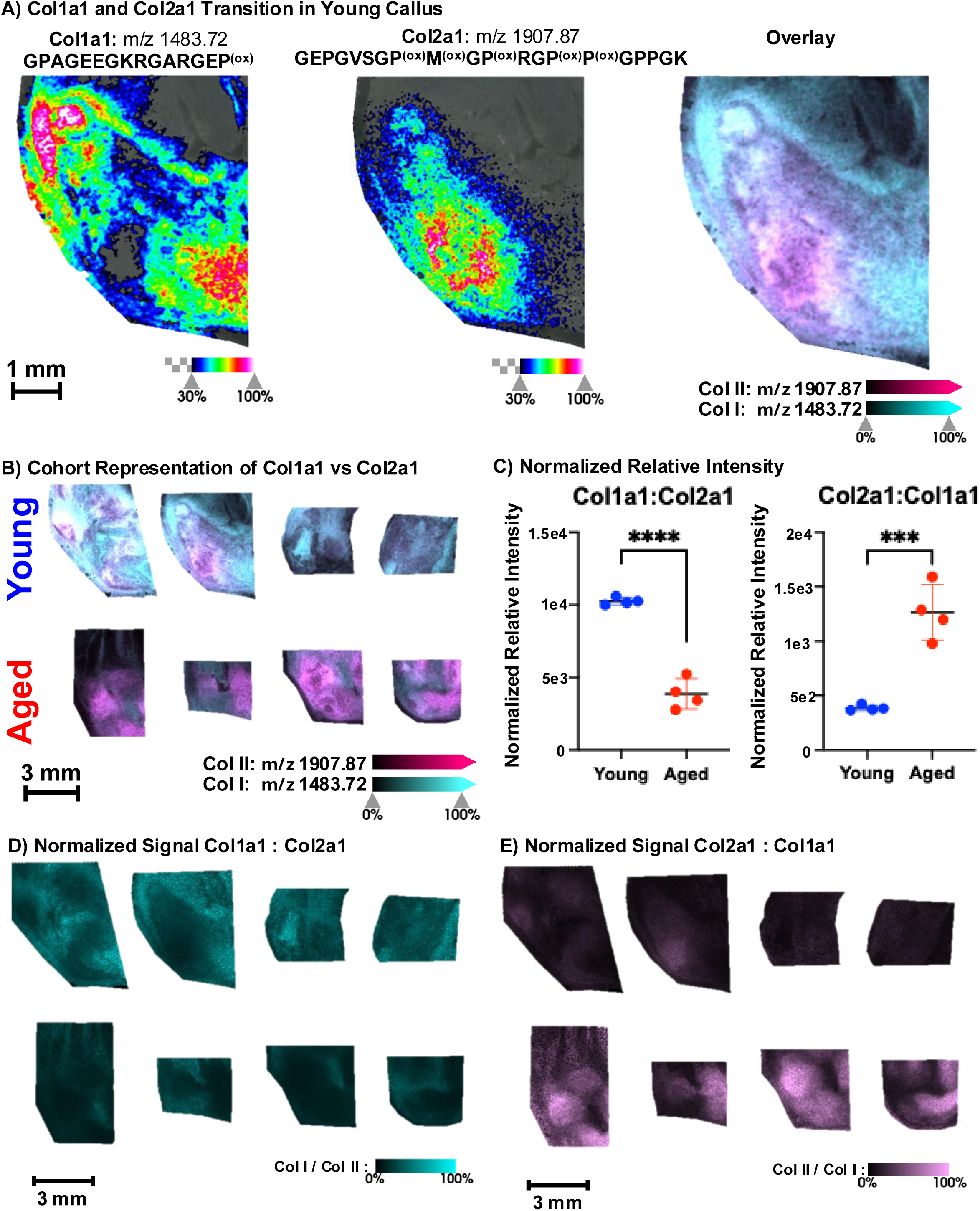
Tracking Osteochondral Remodeling and Delayed Fracture Remodeling with Age. **(A)** Heatmap of signal in a young representative sample from identified peptides of the bone marker protein Col1a1, G^450^PAGEEGKRGARGEP^(ox)-464^, at m/z 1483.72 and the cartilage marker protein Col2a1, G^513^NRGFP^(ox)^GQDGLA^524^, at m/z 1204.56. **B)** Overlay of these two peptide signals shows the transition of matrix proteins and tissue types as the callus turns over from cartilage (pink) to bone (blue) with almost discrete separation in areas of highest signal from the two marker peptides. Visualizing these peptides in all samples **(C)** shows more areas of defined areas of blue, representing bone matrix, in the young samples (top) than in aged fracture calluses (below). Numerically comparing the relative intensity of the two peptides **(D)**, COL1A1 at m/z 1483.72 and COL2A1 at m/z 1204.56, in young tissue shows that the young callus trends toward upregulating the COL1A1 peptide and repressing the COL2A1 peptide as callus remodeling progresses. Aged tissue does not upregulate COL1A1 and COL2A1 signal remains significantly higher than COL1A1. *p<0.05

Given this initial visualization depicted tissue of a young mouse, we sought to determine if this transition between Col1a1 and Col2a1 was conserved in the callus sections of the older mice. When depicted across all 8 tissue samples (**Figure 4C**), it was clear that the abundance of Col1a1 (blue) was higher in the younger samples while the peptide of Col2a1 (pink) had stronger relative abundance in the aged tissues than for Col1a1, indicating a delay in endochondral ossification in tissue from aged mice. Quantification of the abundance of each peptide via normalized signal intensity in each respective tissue section (**Figure 4D**) determined that the normalized signal for Col2a1 was significantly higher in the aged tissues than the normalized intensity for Col1a1 in the same tissues (p = 0.0299 in a Students t-test). Interestingly, normalized signal intensity for Col1a1 in the young samples only trended (p = 0.0749) toward upregulation compared to the normalized signal of Col2a1. This perhaps indicated that the emergence of Col1a1 and the loss of Col2a1 are not directly proportional – but still linked – during fracture healing and that Col2a1 and cartilage were still present in the healing young callus at 10 days post fracture.

Detection of peptides for Col2a1 in callus sections from young animals, in contrast to our earlier finding that other peptides for Col2a1 were significant predictors for aged tissue sections only. This was likely due to the broad classification of experimental groups solely by age, rather than completing a more tissue specific comparison that would identify regions of tissue across the age groups that had more similar molecular identity. To investigate more granular changes in tissue type regardless of age, automated region segmentation was performed to generate an Unbiased Hierarchical Clustering Tissue Segmentation across all tissues simultaneously (**Figure 5A, Supplemental Figure 5A,B**). This process identified four major tissue regions across all eight imaged fracture calluses. Spectra from the two largest regions, Green (149,734 pixels) and Yellow (85,759 pixels) were selected for downstream analysis. The Green ROI was selected due to its predominant distribution among the young tissue samples and its resemblance to the areas of bony callus from H&E staining, and was termed “Bone-like”, while the Yellow ROI showed a more proportional area within the tissues from aged mice or for areas of known cartilaginous tissues by H&E staining and was termed “Cartilage-like”. Portions of the “Bone-like” and “Cartilage-like” regions were still present across both age groups. The next largest identified region, Pink (56,872 pixels), visually appeared to match areas of muscle or soft tissue shown by H&E staining and was not considered for analysis of the fracture callus. A second Discriminating Feature analysis (AUROC >0.7) was completed to find features that could distinguish the newly annotated tissue regions “Bone-like” and “Cartilage-like” in a tissue specific, age-agnostic manner. 43 unique m/z candidate features were found with predictive AUROC values for either the “Bone-like” and “Cartilage-like” regions (**Supplemental Table 1**). These 43 features included 21 new features not found in the comparison for Young vs Aged tissue regions (**Figure 2**). This is likely due to the more sophisticated tissue segmentation operation which used the spatial distribution as the key delineator rather than age. For instance, a newly identified peptide derived from Elastin (Eln m/z 826.45 G^340^GAGAIPGIGG^350^) **Supplemental Figure 5C**) was highly abundant in the aged tissue samples but was also still detectable in the young callus group. Thus, the AUROC score for predicting young vs aged groups based on this ion was below significance (Young vs Aged regions AUROC = 0.468, |AUROC| = 0.532). However, when evaluating this peptide of Elastin for its predictive ability for the newly segmented “Bone-like” and “Cartilage-like” ROIs, the AUROC value achieved statistical significance for predicting the “Cartilage-like” ROI (“Bone-like” and “Cartilage-like” regions AUROC = 0.154, |AUROC| = 0.846).

**Figure 5:**
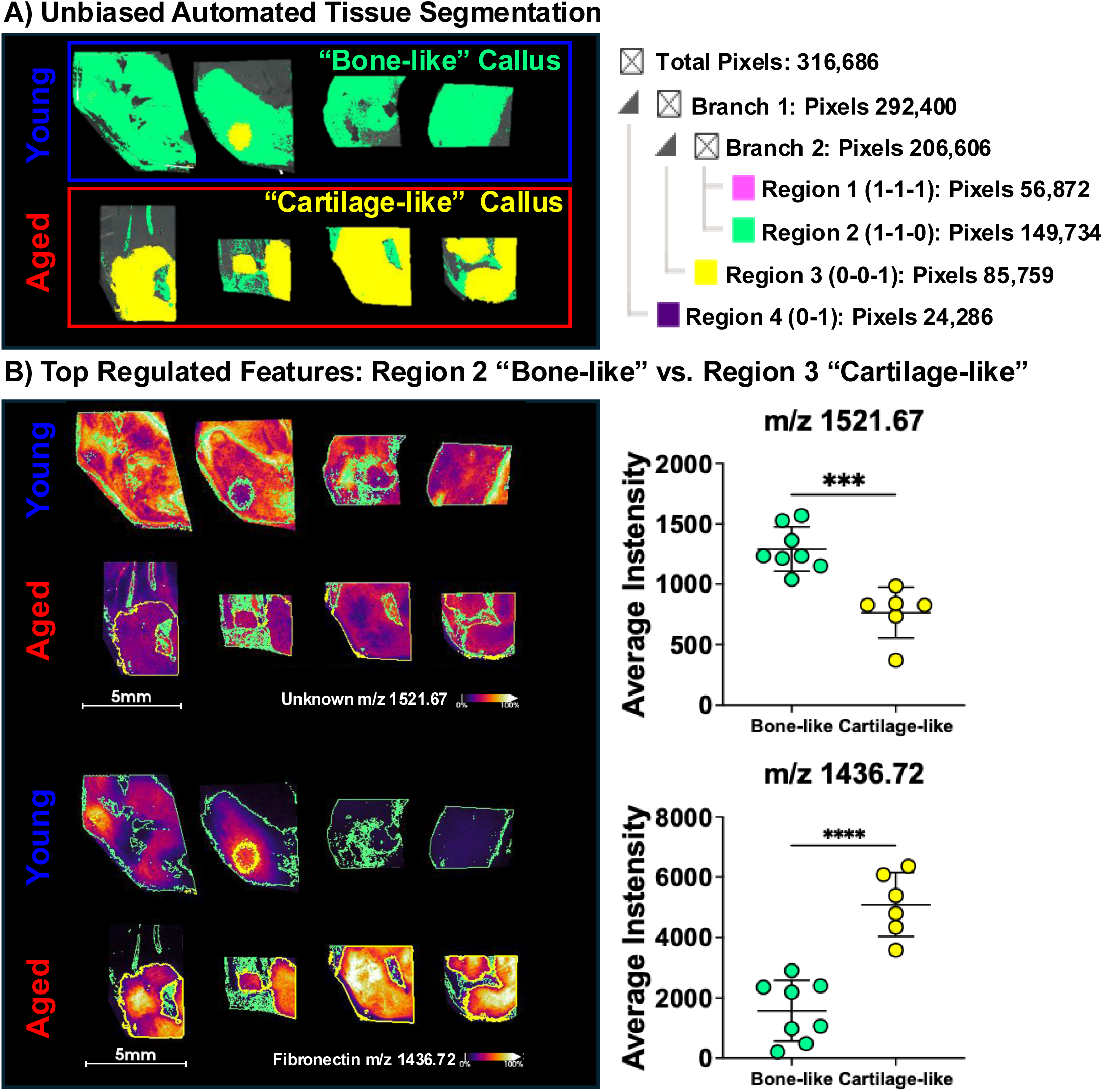
Unbiased Tissue Segmentation by Molecular Signature Increases Feature Identification. **A)** Automated region segmentation on using a Bisecting K-means method utilizing Manhattan Distance was applied to the top 500 most abundant peaks from the composite spectrum. This resulted in several tissue regions across both age groups that were similar to each other in molecular identity. The two largest of these segments, Region 2 (Green 149,734 spectra/pixels) and Region 3 (Yellow 85,759 spectra/pixels) matched identified tissue types from H&E staining and were arbitrarily termed “Bone-like” and “Cartilage-like” tissue regions, respectively. Discriminating feature analysis utilizing the ROC was performed using the new tissue regions as classes and identified 43 features with a predictive |AUROC| > 0.7. **B)** Spatial Heatmaps for molecular features with significant regulation by tissue type alone (“Bone-like” vs “Cartilage-like”) are shown for two of the most significantly regulated features, an unknown peptide at m/z 1521.67 with higher abundance in the “Bone-like” region and a peptide of Fibronectin, F^1584^TVPGSKSTATINN^1597^ at m/z 1436.72 with higher abundance in the “Cartilage-like” region. Green and Yellow outlines in the figure depict the borders of each identified tissue region.

In order to validate the new set of candidate features individually across the samples, the average pixel intensities were isolated by the new tissue sub-regions that were annotated by both tissue age and tissue region (**Figure 5A**). The average intensity for each candidate m/z from the new automatically generated sub-regions, “Bone-like” and “Cartilage-like”, was quantitatively compared with a paired student’s t-test. Spatial heatmaps for two identified candidate m/z features are shown (**Figure 5B**) for a candidate m/z’s including an unknown feature m/z 1521.67 and an identified peptide derived from Fibronectin (Fn) m/z 1436.72 (F^1584^TVPGSKSTATINN^1597^). Quantification and comparison of their abundance between the “Bone-like” (green outlines) and “Cartilage-like” (yellow outlines) regions found that abundance for m/z 1521.67 was significantly higher in the “Bone-like” ROIs while the Fibronectin peptide at m/z 1436.72 was statistically more abundant in the “Cartilage-like” ROIs. Statistical analysis for the complete 43 candidate features identified through Discriminating Feature Analysis via a paired t-test resulted in 27 features retaining a significant difference in ion abundance between the “Bone-like” and “Cartilage-like” ROIs (**Supplemental Figure 6**).

Hierarchical Clustering of feature intensity by subregion revealed a further level of nuance to the regulation of molecular features across the different callus tissues (**Figure 6A**). Each tissue subregion primarily retained its earlier separation first by age group, with subregions belonging to the “Young” group clustering on the left and tissues from the “Aged” group clustered on the right in the hierarchical heatmap. However, when observing the bone and cartilage-like annotations, “Young, Cartilage-like” tissues and “Age, Bone-like” subregions belonged to an ‘intermediate’ branch in the cluster dendrogram and defined a third tissue class between the two extremes (“Young, Bone-like”, “Aged, Cartilage-like”). Re-classifying the spatial ROI’s as these three new stages (**Figure 6B**) as I - Pink, II – Orange, and III - Blue allowed for another more detailed layer of assessment of the fracture healing processes that segmented tissues according to the progression of healing from the earliest, cartilaginous callus (Blue) onward to the more later, harder boney callus (pink). Following spatial region re-classification, one-way ANOVA utilizing each new class (I, II, III) as factors was used to determine the regulation of each feature across the new gradient-defined tissue regions. In this analysis 36 of the 43 features identified showed a significant difference in this manner, revealing more detailed spatial regulation than the binary “Bone-like” vs “Cartilage-like” classification alone that only showed 27 features with statistical differences (**Supplemental Figure 7**). The heatmaps display features that showed significant regulation with the ANOVA-tissue gradient classification system, which however did not show significant regulation in the binary region classification of “Bone-like” vs “Cartilage-like”, m/z 1855.91 and m/z 912.52, (**Figure 6C**). This illustrates the effectiveness and importance of understanding molecular regulation of matrix material at increasingly finer resolution than would be possible with traditional histological techniques. More examples of identified molecular features that have significant regulation by the tissue gradient classification, including an additional peptide for Col2a1, and identified peptides for Elastin and Fibronectin are depicted in **Supplemental Figure 8**.

**Figure 6:**
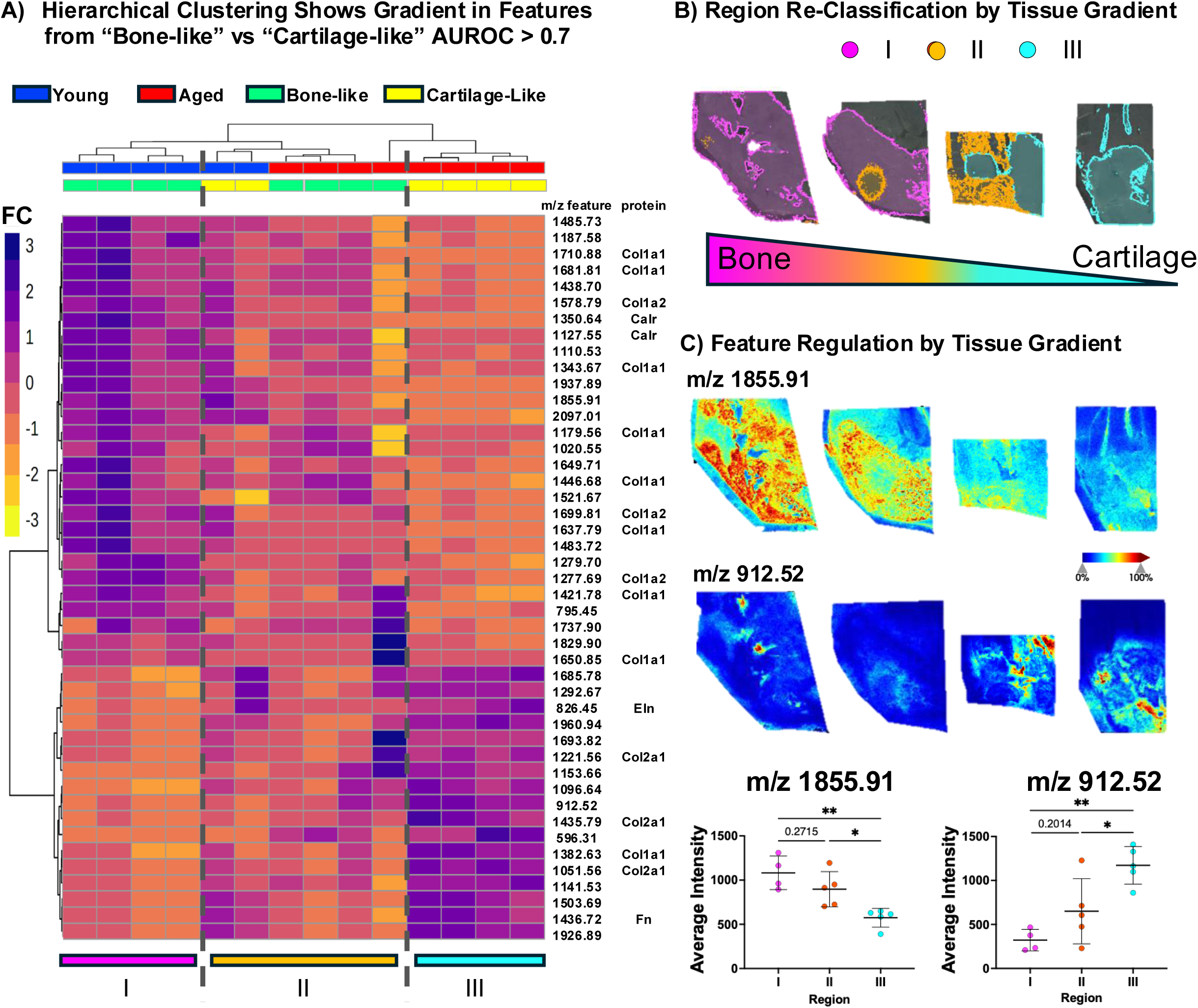
Hierarchical Clustering of Feature Intensities by Sub-region Reveals Gradient of Tissue Healing Across Age Groups. **A)** Clustering the intensities for features identified by Discriminating Feature Analysis between the “Healed” and “Delayed” tissue regions with annotations for each sub-regions Age and Tissue type tissue type showed separation by age and a mixing of tissue types. Notably, though age groups separated in hierarchical clustering, aged-healed regions clustered closer to young-delayed regions than the rest of the aged-delayed samples. Visualized in the heatmap, this resulted in an internal “transitional” set of tissues revealing a gradient of tissue types throughout the healing process. **B)** Using the three regions depicted by the heatmap, tissue boundaries were redrawn and signal reexported using these annotations. **C)** Visual representation of selected features at m/z 1187.58 and 1960.94 illustrate the gradient of change that occurs that is not entirely dictated by age of the tissue. ANOVA was completed on exported feature intensities using the three tissue regions as factors with Bonferroni corrected pair-wise post hoc comparisons to illustrate the progressive change of the molecular feature throughout the healing process across young and aged tissues. *p<0.017, **p<0.003, ***p<0.0003 in 3-way Bonferroni corrected *post hoc* pairwise comparison after ANOVA.

## Discussion

Age-related delays in fracture healing are common and contribute substantially to morbidity and mortality in the elderly^1,2^. Clinical and animal studies demonstrate slower healing with delayed cartilage and bone formation, impaired resorption, and an increased risk of delayed union or non-union requiring surgical intervention^7–14^. Although healing time varies with comorbidities, delayed fracture repair remains a major clinical challenge, highlighting an unmet need for therapies that accelerate healing in an aging population. Identifying molecular regulators of fracture repair is essential to develop interventions that reduce morbidity and mortality. Matrix-Assisted Laser Desorption/Ionization Mass Spectrometry Imaging (MALDI MSI) enables spatially resolved, in situ analysis of proteins, lipids, metabolites, and glycans directly from tissue sections situ^37–41^. Widely applied to cancer and neurodegenerative diseases, spatial MS Imaging has more recently been used to study musculoskeletal tissues^46–57,68^. Recently, we utilized a collagenase type III digestion on human knee tissue, enabling spatial proteomics of bone-cartilage composite tissues for analysis of highly crosslinked bone and cartilage ECM^62^. This builds on the growing number of applications for the use of MS Imaging for multi-spatial-omics for musculoskeletal (MSK) tissues. We have previously completed a standard analysis of healing rates in animals of different ages^9^, and using materials from that study^9^, we applied the collagenase-assisted spatial ECM proteomics to bone–cartilage tissues in mice for the first time to identify key molecular checkpoints underlying age-related delays in fracture healing.

We present the first protein-based spatial molecular mapping of murine skeletal tissues undergoing endochondral ossification by targeting extracellular matrix proteins via collagenase III digestion. Spatial MS Imaging of enzyme-digested tissue generated feature-rich spectra that clearly distinguish bone from cartilage in the callus sections from young (3-month-old) and aged (18-month-old) animals. This emphasizes the impact and utility of this approach for interrogating musculoskeletal disease and ECM complexity in model organisms. Detected proteins included canonical tissue markers, such as Col1a1 in bone and Col2a1 in cartilage, with spatial localization matching the areas of identified bone and cartilage tissues via H&E staining. Unbiased molecular image processing enabled simultaneous analysis of hundreds of peptide features, allowing for a molecular visualization and mapping of the callus healing process. At 10 days post tibial fracture, MS Imaging identified increased markers of cartilaginous tissues in the aged callus tissue sections, identifying markers of delayed healing through matrix markers. For instance, all identified peptides for Col2a1 were significantly more abundant in aged tissue sections than in young callus sections. Conversely, several candidate marker peptides with a significantly predictive AUROC score for young tissue were peptides of collagen type I, a protein normally associated with bone matrix. Simultaneous visualization for peptides of Col1a1 and Col2a1 in the same tissues allowed for direct assessment of the progression of healing in young vs aged callus sections, identifying that at 10 days post-fracture, endochondral ossification was significantly delayed in the aged fracture callus.

The specificity of MS Imaging for identifying peptide markers in place on tissue increases the ability to delineate tissue types in complex, composite tissues such as the fracture callus. Using an unbiased, label-free hierarchical segmentation, young and aged callus tissues were segmented into several classes or regions of tissue with molecular similarity. This revealed that while in general endochondral ossification had occurred to a greater extent in the younger animals 10 days after injury compared to aged animals, this process is highly dynamic and locally regulated, and cartilage was still present in areas of the callus from young animals. The molecular signature of the aged tissues, where cartilage tissues are more abundant, can be used to identify cartilage tissues in young animals where remodeling is incomplete. Thus, spatial MS Imaging can identify target regions of dynamic tissue remodeling that may be targeted for follow-up studies investigating matrix dynamics on even shorter length scales. While this example provides only a snapshot of the dynamic process of matrix remodeling, future longitudinal works may be able to utilize these tissue signatures to track and assess tissue healing in response to interventions. Further, new insights from spatial MS Imaging included the identification of important intermediate steps in the healing process. For instance, areas of “bone-like” tissue in the aged callus sections more molecularly resembled the “cartilage-like” tissues from young samples, as indicated by hierarchical clustering by tissue type in **Figure 6A**. This implies that even bone-like fracture tissue generated by aged-animals, identified via H&E in other standard approaches, is molecularly distinct from the bone-like tissues generated by younger callus tissues at the same time. This raises important clinical considerations about how aged-fracture patients should be addressed, as the aged tissues perhaps fail to overcome molecular hurdles or checkpoints easily surpassed in young contexts that would allow the formation of more mature bone-like tissues. This “intermediate” tissue composed of a mix of both bone and cartilage-like materials is easily identified via MADLI MSI and may be missed in other histopathological imaging approaches.

Beyond simple tracking of marker peptides and proteins, spatial MS Imaging provided novel biological insights into tissue progression that occurs during the fracture healing process and which may contribute to delayed healing. For instance, the matrix protein fibronectin (Fn) was identified as characteristic of the “cartilage-like” or least matured areas of callus tissue in the aged-samples. Fn has several known roles in fracture healing most notably associated with the formation of the initial hematoma following immediate injury^69^. Importantly, Fn can act to form an “emergency” ECM following tissue damage and can bind and house important signaling factors that regulate bone formation, including TGFb among others^70–72^. While FN may be expressed by chondrocytes and osteoblastic cells within the callus and contribute to bone healing^73^, it is also expressed by neutrophils that infiltrate the fracture hematoma at the time of injury^72^. Neutrophils are key regulators and drivers of the phenomena known as “inflammaging”, the chronic state of inflammation associated with aging that can drive tissue damage and promote chronic disease and frailty^74^. Thus, the prolonged presence of Fn in the aged callus raises concerns about the mechanisms of fracture healing in aged tissues: Fn may simply be a bystander of prolonged resolution towards osteochondral tissues or it may be the result of an active process of inflammatory cell types delaying the progression of local healing. Future works targeting the role of Fn in fracture healing may inform future treatments for delayed fracture healing in aged patients. Overall, Fn in the callus from the aged tissue sections indicates a much earlier stage of callus remodeling or delayed healing. Similarly, the basement protein Elastin (Eln) was also found upregulated in the “cartilage-like” aged fracture callus tissues. Eln has previously been identified as a marker for the early soft-callus that forms quickly after injury, indicating production of connective tissue proteins that are later remodeled into the mineralized callus^75^. Presence of Fn and Eln in the aged, “cartilage-like” callus regions indicates that healing by 10-days post fracture in the 18-month-old mice progresses slower than the 3-month-old animals.

Novel functions were found by spatial MS Imaging in the young fracture callus that may promote healthy healing responses. Multiple peptides for Calreticulin (Calr) were detected that had a significant up-regulation only in the young and “bone-like” callus tissues. Calr is a highly versatile calcium-binding chaperone with multiple known functions in the endoplasmic reticulum (ER) participating in the synthesis of a variety of molecules, including ion channels, surface receptors, integrins, and transporters. Calr also affects intracellular Ca(^2+^) homoeostasis by modulation of ER Ca(^2+^)^76^. Calr currently has no confirmed roles specifically in fracture healing, but it has been documented with roles in the skeleton regulating bone formation and resorption. For example, Calr treatment showed inhibited inflammatory osteoclastogenesis of RANKL and LPS-treated osteoclast precursor cells through inhibition of c-Fos and nuclear factor of activated T cells, cytoplasmic 1 (Nfatc1)^77^. Perhaps more importantly, and directly related to fracture callus remodeling, Calr in mouse embryonic cells acts as a molecular switch to osteoblastogenesis over chondrogenesis by promoting the intranuclear transport of Nfatc1 and the nuclear translocation of β-catenin required for commitment to osteogenic lineages^78^. In other contexts, expression of Calr may decrease with age^79^, where it would otherwise act to regulate cellular senescence and enhance wound repair^80^. Given the importance of the transition from chondrogenic cells to osteogenic cell types in the turnover from soft to hard callus during fracture healing, and the potential for these functions to be regulated with age, the relevance of Calr in the fracture healing process is intriguing. This detection of multiple peptides of Calreticulin confirms its presence in the fracture callus and offers new insights for a protein that may act as an important “checkpoint” regulating the transition from cartilaginous tissues to mineralized bone in callus healing – a checkpoint often delayed or absent in aged fracture callus healing processes.

We present the utilization of spatial MS Imaging in conjunction with a collagenase type III enzymatic pre-treatment for spatial proteomics on murine FFPE fracture callus tissue slides. Using tissue sections from both young and aged animals, we demonstrate that aged animals exhibit a profound delay in callus remodeling 10 days-post tibial fracture by tracking matrix proteins. Additionally, we visualize the transition between bone and cartilage like tissues in the healing callus to reveal the gradient of transition between soft and card callus materials and depict the absence of this transition in the tissue sections of aged animals. Further we identify other non-matrix proteins, namely Fibronectin and Calreticulin, that may inhibit or promote important transitional steps between cartilage and bone materials critical to the progression of healing in the fracture callus. Further work will determine the functional relevance of these non-matrix proteins in fracture healing with the goal of generating therapies to promote healing in elderly patients. However, direct detection and tracking of matrix remodeling and non-matrix proteins throughout the fracture healing process by spatial MS Imaging demonstrated new innovative approaches in musculoskeletal research.

## Acknowledgements

We acknowledge the NIH for support from the following granting agencies and mechanisms: NIA T32 AG000266 (to Schurman, PI: Ellerby), NIA P01 AG066591 – (PI: Ellerby, Core Director: Schilling), NIAMS R21 AR084303 (MPIs: Schilling, Alliston, Angel) and NIH/OD S10 OD030212 (PI: Angel), NIH/OD S10 OD038264 (PI: Schilling), NCI R21 CA240148 (PI: Angel).

## Figure Captions

**Supplemental Figure 1: MALDI MSI workflow.** FFPE tissue slides are prepared for Matrix Assisted Laser Desorption/Ionization (MALDI) Mass Spectrometry Imaging (MSI) through serial application of PNGase F and Collagenase III followed by application of α-Cyano-4-hydroxycinnamic acid (CHCA) matrix. Single pixel mass spectra are collected from prepared tissue sections with the timsTOF fleX (Bruker). Collected spectra are combined to generate the mean MALDI MSI spectra where individual peaks, or m/z features, can be isolated to generate spatial heatmaps of molecular features across imaged tissue. To assign molecular identifications to the m/z features, imaged tissues are reclaimed from the slide and prepared for LC-MS/MS analysis utilizing the same enzymes that prepared the tissues for imaging.

**Supplemental Figure 2: Identification of Fracture Callus Region.** Hematoxylin and Eosin (pink, left) histologic staining and unstained, enzymatically digested tissue sections (right, grey) of unfixed fracture calluses from (A) young (3 mo.) and (B) aged (18 mo.) mice 10 weeks post tibial fracture showing regions of tissue selected for MALDI MSI (black dotted box, left or red solid region, right).

**Supplemental Figure 3: Discriminating Features Analysis by Age**: SCiLS lab identified 31 unique m/z features with an |AUROC| > 0.7 that had significant predictive ability for either the young or aged fracture callus. Exported intensities for each feature from each tissue section are compared showing cohort level significant difference in a t-test for 29 of 31 features demonstrating the consistency of regulation of each feature by age. The remaining two features trended toward significance with a p value <0.1. *p<0.05 **p<0.01 ***p<0.001

**Supplemental Figure 4: Spatial Distribution for Calreticulin as regulated by Age in Fracture Callus.** Ion intensity heatmaps for the two peptides of Calreticulin, (A) V^321^KSGTIFDNF^330^ m/z =1127.56 and (B) T^288^WIHPEIDNPE^298^ m/z = 1350.64 were identified as upregulated in the young callus group and largely shared a similar spatial distribution, further indicating each unique peptide stemmed from the same protein.

**Supplemental Figure 5: Tissue Classification by Automated Segmentation.** A) Hierarchical segmentation based on molecular signal from MALDI MSI on fracture callus tissue samples identified several distinct tissue types. The two largest regions matched areas from H&E staining that identified more “bone-like” (Region 2, green) and “cartilage-like” (Region 3, yellow) tissues areas across both young and aged callus groups. B) Sub-region separation using the dual labels of age-group (young, aged) and tissue type (bone or cartilage-like) resulted in 14 unique ROIs for feature extraction. C) Utilizing the tissue-specific ROIs, instead of age-group alone, resulted in higher AUROC predictive scores for several peptides increasing the number of identified candidates for validation, as exhibited by the identified peptide for Elastin G^340^GAGAIPGIGG^350^ at m/z 826.45 that had an AUROC value increase from a non-significant/non-predictive score by age of AUROC = 0.468 to a significant AUROC = 0.154 (|AUROC| = 1 - 0.154 = 0.846) when using the tissue segmentation.

**Supplemental Figure 6: Discriminating Features Analysis by Tissue Type**: SCiLS lab identified 43 unique m/z features with an |AUROC| > 0.7 that had significant predictive ability for the automatically generated tissue regions termed “Bone-like” (Fig. 5B, green) and “Cartilage-like” (Fig. 5B, yellow). Exported intensities for each feature from each of the 14 identified tissue sub-regions are compared showing cohort level significant difference in a paired t-test for 27 of 43 features. Of the remaining features, 11 features trended toward significance with a p value <0.1. *p<0.05 **p<0.01 ***p<0.001

**Supplemental Figure 7: Regulation of Identified Features after Gradient Re-classification:** After hierarchical clustering sorted the 14 exported tissue sub-regions subregions into 3 new tissue classes showing a gradient of healing/matrix remodeling throughout fracture healing, 36 of the 43 features found by Discriminating Feature analysis showed a significant difference in one-way ANOVA when utilizing each new class (I, II, III) as factors. 3-way Bonferroni corrected post-hoc pairwise comparisons are shown for features that showed significant difference in initial ANOVA testing. *p<0.017, **p<0.003, ***p<0.0003, ****p<0.0001.

**Supplemental Figure 8: Select Ion Heatmaps for Features Identified by MALDI MSI with Tissue Type Regulation.** Ion intensity heatmaps for three peptides identified with differential abundance across different tissue regions (A) within the fracture callus. B) Collagen II peptide G^1101^PAGARGIAGPQ^1112^ m/z = 1051.56, C) Elastin peptide G^340^GAGAIPGIGG^350^ m/z 826.45 and D) Fibronectin peptide F^1584^TVPGSKSTATINN^1597^ m/z 1436.72 show higher abundance in the more cartilaginous (III) region of the callus compared to the intermediate (II) or more boney (I) regions.

## Notes

### Competing Interest Statement

Dr. Nannan Tao is employee of Bruker.

### Funding

- National Institutes of Health: S10OD038264, R21AR084303, T32AG000266, P01AG066591, S10OD030212, R21CA240148

## Citations

1 Cauley, J. A., Thompson, D. E., Ensrud, K. C., Scott, J. C. & Black, D. Risk of mortality following clinical fractures. Osteoporos Int 11, 556–561 (2000).

2 White, B. L., Fisher, W. D. & Laurin, C. A. Rate of mortality for elderly patients after fracture of the hip in the 1980’s. J Bone Joint Surg Am 69, 1335–1340 (1987).

3 ElHawary, H. et al. Bone Healing and Inflammation: Principles of Fracture and Repair. Semin Plast Surg 35, 198–203 (2021).

4 Geerts, W. H. et al. Prevention of venous thromboembolism. Chest 119, 132S–175S (2001).

5 Rose, S. & Maffulli, N. Hip fractures. An epidemiological review. Bull Hosp Jt Dis 58, 197–201 (1999).

6 Ghimire, S. et al. The investigation of bone fracture healing under intramembranous and endochondral ossification. Bone Rep 14, 100740 (2021).

7 Molitoris, K. H., Balu, A. R., Huang, M. & Baht, G. S. The impact of age and sex on the inflammatory response during bone fracture healing. JBMR Plus 8, ziae023 (2024).

8 Clark, D., Nakamura, M., Miclau, T. & Marcucio, R. Effects of Aging on Fracture Healing. Curr Osteoporos Rep 15, 601–608 (2017).

9 Lu, C. et al. Cellular basis for age-related changes in fracture repair. J Orthop Res 23, 1300–1307 (2005).

10 Meinberg, E. G., Clark, D., Miclau, K. R., Marcucio, R. & Miclau, T. Fracture repair in the elderly: Clinical and experimental considerations. Injury 50 Suppl 1, S62–S65 (2019).

11 Claes, L. et al. Monitoring and healing analysis of 100 tibial shaft fractures. Langenbecks Arch Surg 387, 146–152 (2002).

12 Hee, H. T., Wong, H. P., Low, Y. P. & Myers, L. Predictors of outcome of floating knee injuries in adults: 89 patients followed for 2-12 years. Acta Orthop Scand 72, 385–394 (2001).

13 Aho, A. J. Electron microscopic and histologic studies on fracture repair in old and young rats. Acta Chir Scand Suppl 357, 162–165 (1966).

14 Lopas, L. A. et al. Fractures in geriatric mice show decreased callus expansion and bone volume. Clin Orthop Relat Res 472, 3523–3532 (2014).

15 Foulke, B. A., Kendal, A. R., Murray, D. W. & Pandit, H. Fracture healing in the elderly: A review. Maturitas 92, 49–55 (2016).

16 Naik, A. A. et al. Reduced COX-2 expression in aged mice is associated with impaired fracture healing. J Bone Miner Res 24, 251–264 (2009).

17 Yokoyama, K. et al. Evaluation of functional outcome of the floating knee injury using multivariate analysis. Arch Orthop Trauma Surg 122, 432–435 (2002).

18 Hernandez, R. K., Do, T. P., Critchlow, C. W., Dent, R. E. & Jick, S. S. Patient-related risk factors for fracture-healing complications in the United Kingdom General Practice Research Database. Acta Orthop 83, 653–660 (2012).

19 Xia, S. et al. An Update on Inflamm-Aging: Mechanisms, Prevention, and Treatment. J Immunol Res 2016, 8426874 (2016).

20 Josephson, A. M. et al. Age-related inflammation triggers skeletal stem/progenitor cell dysfunction. Proc Natl Acad Sci U S A 116, 6995–7004 (2019).

21 Hofbauer, L. C., Baschant, U. & Hofbauer, C. Targeting senescent cells to boost bone fracture healing. J Clin Invest 134 (2024).

22 Saul, D. et al. Modulation of fracture healing by the transient accumulation of senescent cells. Elife 10 (2021).

23 Bucher, C. H. et al. Local immune cell contributions to fracture healing in aged individuals - A novel role for interleukin 22. Exp Mol Med 54, 1262–1276 (2022).

24 Clark, D. et al. Age-related changes to macrophages are detrimental to fracture healing in mice. Aging Cell 19, e13112 (2020).

25 Lin, T. H. et al. Decreased osteogenesis in mesenchymal stem cells derived from the aged mouse is associated with enhanced NF-kappaB activity. J Orthop Res 35, 281–288 (2017).

26 Reikeras, O., Shegarfi, H., Wang, J. E. & Utvag, S. E. Lipopolysaccharide impairs fracture healing: an experimental study in rats. Acta Orthop 76, 749–753 (2005).

27 Xing, Z., Lu, C., Hu, D., Miclau, T., 3rd & Marcucio, R. S. Rejuvenation of the inflammatory system stimulates fracture repair in aged mice. J Orthop Res 28, 1000–1006 (2010).

28 Baht, G. S. et al. Exposure to a youthful circulaton rejuvenates bone repair through modulation of beta-catenin. Nat Commun 6, 7131 (2015).

29 Meyer, R. A., Jr., et al. Gene expression in older rats with delayed union of femoral fractures. J Bone Joint Surg Am 85, 1243–1254 (2003).

30 Mancinelli, L. & Intini, G. Age-associated declining of the regeneration potential of skeletal stem/progenitor cells. Front Physiol 14, 1087254 (2023).

31 Vi, L. et al. Macrophage cells secrete factors including LRP1 that orchestrate the rejuvenation of bone repair in mice. Nat Commun 9, 5191 (2018).

32 Clark, D. et al. Age-related changes to macrophage subpopulations and TREM2 dysregulation characterize attenuated fracture healing in old mice. Aging Cell 23, e14212 (2024).

33 Lu, C. et al. Effect of age on vascularization during fracture repair. J Orthop Res 26, 1384–1389 (2008).

34 Meyer, R. A., Jr., et al. Age and ovariectomy impair both the normalization of mechanical properties and the accretion of mineral by the fracture callus in rats. J Orthop Res 19, 428–435 (2001).

35 Hillenkamp, F., Karas, M., Beavis, R. C. & Chait, B. T. Matrix-assisted laser desorption/ionization mass spectrometry of biopolymers. Anal Chem 63, 1193A–1203A (1991).

36 Nakanishi, T., Okamoto, N., Tanaka, K. & Shimizu, A. Laser desorption time-of-flight mass spectrometric analysis of transferrin precipitated with antiserum: a unique simple method to identify molecular weight variants. Biol Mass Spectrom 23, 230–233 (1994).

37 Franck, J. et al. MALDI imaging mass spectrometry: state of the art technology in clinical proteomics. Mol Cell Proteomics 8, 2023–2033 (2009).

38 Walch, A., Rauser, S., Deininger, S. O. & Hofler, H. MALDI imaging mass spectrometry for direct tissue analysis: a new frontier for molecular histology. Histochem Cell Biol 130, 421–434 (2008).

39 Heijs, B. et al. Molecular signatures of tumor progression in myxoid liposarcoma identified by N-glycan mass spectrometry imaging. Lab Invest 100, 1252–1261 (2020).

40 Angel, P. M., Mehta, A., Norris-Caneda, K. & Drake, R. R. MALDI Imaging Mass Spectrometry of N-glycans and Tryptic Peptides from the Same Formalin-Fixed, Paraffin-Embedded Tissue Section. Methods Mol Biol 1788, 225–241 (2018).

41 Tuck, M., Grelard, F., Blanc, L. & Desbenoit, N. MALDI-MSI Towards Multimodal Imaging: Challenges and Perspectives. Front Chem 10, 904688 (2022).

42 Michno, W., Wehrli, P. M., Blennow, K., Zetterberg, H. & Hanrieder, J. Molecular imaging mass spectrometry for probing protein dynamics in neurodegenerative disease pathology. J Neurochem 151, 488–506 (2019).

43 Hanrieder, J., Ljungdahl, A. & Andersson, M. MALDI imaging mass spectrometry of neuropeptides in Parkinson’s disease. J Vis Exp (2012).

44 Angel, P. M. et al. Extracellular Matrix Imaging of Breast Tissue Pathologies by MALDI-Imaging Mass Spectrometry. Proteomics Clin Appl 13, e1700152 (2019).

45 Angel, P. M. et al. Zonal regulation of collagen-type proteins and posttranslational modifications in prostatic benign and cancer tissues by imaging mass spectrometry. Prostate 80, 1071–1086 (2020).

46 Kriegsmann, M. et al. MALDI MS imaging as a powerful tool for investigating synovial tissue. Scand J Rheumatol 41, 305–309 (2012).

47 Cillero-Pastor, B., Eijkel, G. B., Blanco, F. J. & Heeren, R. M. A. Protein classification and distribution in osteoarthritic human synovial tissue by matrix-assisted laser desorption ionization mass spectrometry imaging. Anal Bioanal Chem 407, 2213–2222 (2015).

48 Lee, Y. R. et al. Mass Spectrometry Imaging as a Potential Tool to Investigate Human Osteoarthritis at the Tissue Level. Int J Mol Sci 21 (2020).

49 Rocha, B. et al. Identification of a distinct lipidomic profile in the osteoarthritic synovial membrane by mass spectrometry imaging. Osteoarthritis Cartilage 29, 750–761 (2021).

50 Eveque-Mourroux, M. R. et al. Spatially resolved endogenous improved metabolite detection in human osteoarthritis cartilage by matrix assisted laser desorption ionization mass spectrometry imaging. Analyst 144, 5953–5958 (2019).

51 Haartmans, M. J. J. et al. Matrix-assisted laser desorption/ionization mass spectrometry imaging (MALDI-MSI) reveals potential lipid markers between infrapatellar fat pad biopsies of osteoarthritis and cartilage defect patients. Anal Bioanal Chem 415, 5997–6007 (2023).

52 Schaepe, K. et al. Imaging of Lipids in Native Human Bone Sections Using TOF-Secondary Ion Mass Spectrometry, Atmospheric Pressure Scanning Microprobe Matrix-Assisted Laser Desorption/Ionization Orbitrap Mass Spectrometry, and Orbitrap-Secondary Ion Mass Spectrometry. Anal Chem 90, 8856–8864 (2018).

53 Seeley, E. H. et al. Co-registration of multi-modality imaging allows for comprehensive analysis of tumor-induced bone disease. Bone 61, 208–216 (2014).

54 Fujino, Y., Minamizaki, T., Yoshioka, H., Okada, M. & Yoshiko, Y. Imaging and mapping of mouse bone using MALDI-imaging mass spectrometry. Bone Rep 5, 280–285 (2016).

55 Svirkova, A., Turyanskaya, A., Perneczky, L., Streli, C. & Marchetti-Deschmann, M. Multimodal imaging of undecalcified tissue sections by MALDI MS and muXRF. Analyst 143, 2587–2595 (2018).

56 Lee, Y. R. et al. High-Resolution N-Glycan MALDI Mass Spectrometry Imaging of Subchondral Bone Tissue Microarrays in Patients with Knee Osteoarthritis. Anal Chem 95, 12640–12647 (2023).

57 Briggs, M. T. et al. MALDI mass spectrometry imaging of N-glycans on tibial cartilage and subchondral bone proteins in knee osteoarthritis. Proteomics 16, 1736–1741 (2016).

58 Cillero-Pastor, B., Eijkel, G. B., Kiss, A., Blanco, F. J. & Heeren, R. M. Matrix-assisted laser desorption ionization-imaging mass spectrometry: a new methodology to study human osteoarthritic cartilage. Arthritis Rheum 65, 710–720 (2013).

59 Rocha, B., Cillero-Pastor, B., Blanco, F. J. & Ruiz-Romero, C. MALDI mass spectrometry imaging in rheumatic diseases. Biochim Biophys Acta Proteins Proteom 1865, 784–794 (2017).

60 Angel, P. M. et al. Extracellular matrix alterations in low-grade lung adenocarcinoma compared with normal lung tissue by imaging mass spectrometry. J Mass Spectrom 55, e4450 (2020).

61 Clift, C. L. et al. Evaluation of Therapeutic Collagen-Based Biomaterials in the Infarcted Mouse Heart by Extracellular Matrix Targeted MALDI Imaging Mass Spectrometry. J Am Soc Mass Spectrom 32, 2746–2754 (2021).

62 Schurman, C. A. et al. Tissue and extracellular matrix remodeling of the subchondral bone during osteoarthritis of knee joints as revealed by spatial mass spectrometry imaging. Bone Res 14, 14 (2026).

63 Thompson, Z., Miclau, T., Hu, D. & Helms, J. A. A model for intramembranous ossification during fracture healing. J Orthop Res 20, 1091–1098 (2002).

64 Clift, C. L., Drake, R. R., Mehta, A. & Angel, P. M. Multiplexed imaging mass spectrometry of the extracellular matrix using serial enzyme digests from formalin-fixed paraffin-embedded tissue sections. Anal Bioanal Chem 413, 2709–2719 (2021).

65 Angel, P. M. et al. Mapping Extracellular Matrix Proteins in Formalin-Fixed, Paraffin-Embedded Tissues by MALDI Imaging Mass Spectrometry. J Proteome Res 17, 635–646 (2018).

66 Angel, P. M. et al. Advances in MALDI imaging mass spectrometry of proteins in cardiac tissue, including the heart valve. Biochim Biophys Acta Proteins Proteom 1865, 927–935 (2017).

67 Macdonald, J. K. et al. Optimization of Collagenase Proteomics for Improved Mass Spectrometry Imaging Peptide Identification. Anal Chem 97, 7672–7681 (2025).

68 Good, C. J. et al. High Spatial Resolution MALDI Imaging Mass Spectrometry of Fresh-Frozen Bone. Anal Chem 94, 3165–3172 (2022).

69 Klavert, J. & van der Eerden, B. C. J. Fibronectin in Fracture Healing: Biological Mechanisms and Regenerative Avenues. Front Bioeng Biotechnol 9, 663357 (2021).

70 Dallas, S. L. et al. Fibronectin regulates latent transforming growth factor-beta (TGF beta) by controlling matrix assembly of latent TGF beta-binding protein-1. J Biol Chem 280, 18871–18880 (2005).

71 Bonewald, L. F. & Dallas, S. L. Role of active and latent transforming growth factor beta in bone formation. J Cell Biochem 55, 350–357 (1994).

72 Bastian, O. W., Koenderman, L., Alblas, J., Leenen, L. P. & Blokhuis, T. J. Neutrophils contribute to fracture healing by synthesizing fibronectin+ extracellular matrix rapidly after injury. Clin Immunol 164, 78–84 (2016).

73 Kilian, O. et al. mRNA expression and protein distribution of fibronectin splice variants and high-molecular weight tenascin-C in different phases of human fracture healing. Calcif Tissue Int 83, 101–111 (2008).

74 Abdullah, G. A., Akpan, A., Phelan, M. M. & Wright, H. L. The complex role of neutrophils in healthy aging, inflammaging, and frailty. J Leukoc Biol 117 (2025).

75 Erickson, C. B. et al. A timeseries analysis of the fracture callus extracellular matrix proteome during bone fracture healing. J Life Sci (Westlake Village) 3, 1–30 (2021).

76 Michalak, M., Corbett, E. F., Mesaeli, N., Nakamura, K. & Opas, M. Calreticulin: one protein, one gene, many functions. Biochem J 344 Pt 2, 281–292 (1999).

77 Fischer, C. R. et al. Calreticulin inhibits inflammation-induced osteoclastogenesis and bone resorption. J Orthop Res 35, 2658–2666 (2017).

78 Pilquil, C., Alvandi, Z. & Opas, M. Calreticulin regulates a switch between osteoblast and chondrocyte lineages derived from murine embryonic stem cells. J Biol Chem 295, 6861–6875 (2020).

79 Zarate, S. M., Huntington, T. E., Bagher, P. & Srinivasan, R. Aging reduces calreticulin expression and alters spontaneous calcium signals in astrocytic endfeet of the mouse dorsolateral striatum. NPJ Aging 9, 5 (2023).

80 Wang, Q., Li, Q. & Wei, N. CALR promotes corneal epithelial cell proliferation and migration through Wnt7a. Mol Biol Rep 52, 714 (2025).

