## Supplementary material for "Age-related Delays in Osteochondral Remodeling of Fracture Healing Illustrated by Mass Spectrometry Imaging": Suppl Figure 1

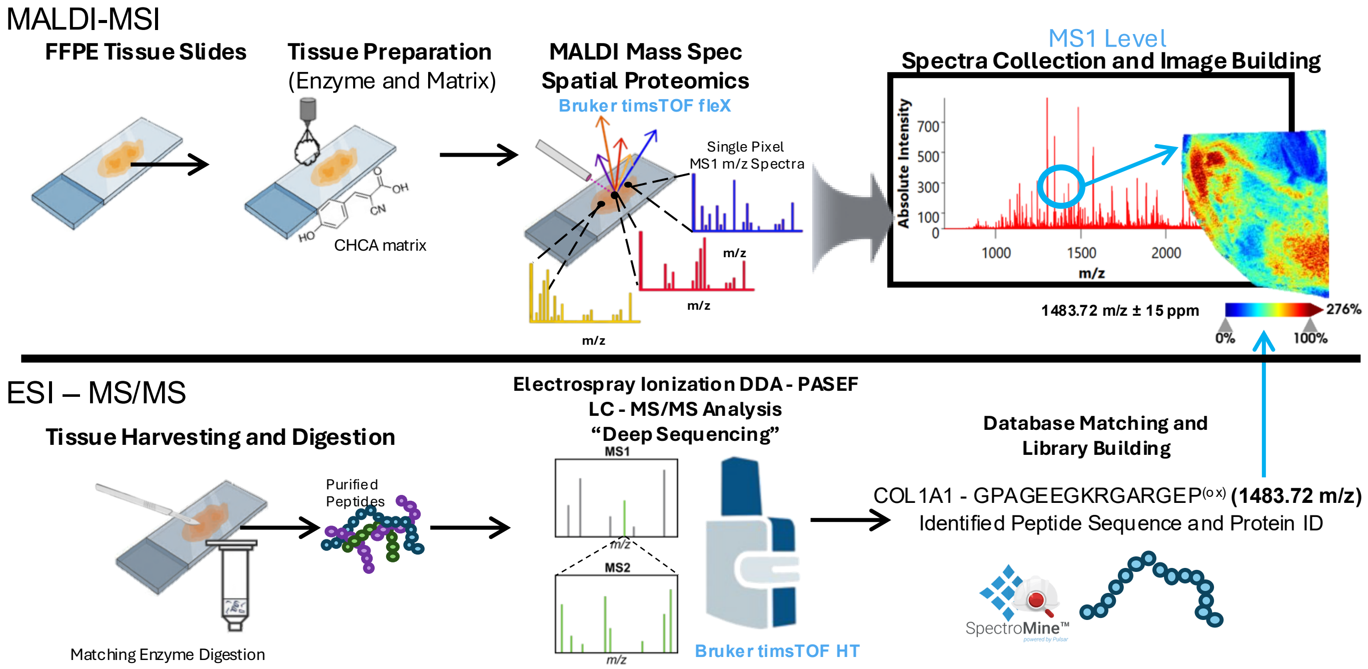


**Supplemental Figure 1: MALDI-MSI workflow.** FFPE tissue slides are prepared for Matrix Assisted Laser Desorption/Ionization (MALDI) Mass Spectrometry Imaging (MSI) through serial application of PNGaseF and Collagenase III followed by application of α-Cyano-4-hydroxycinnamic acid (CHCA) matrix. Single pixel mass spectra are collected from prepared tissue sections with the timsTOF fleX (Bruker). Collected spectra are combined to generate the mean MALDI-MSI spectra where individual peaks, or m/z features, can be isolated to generate spatial heatmaps of molecular features across imaged tissue. To assign molecular identifications to the m/z features, imaged tissues are reclaimed from the slide and prepared for LC-MS/MS analysis utilizing the same enzymes that prepared the tissues for imaging.
