## Supplementary material for "Age-related Delays in Osteochondral Remodeling of Fracture Healing Illustrated by Mass Spectrometry Imaging": Suppl Figure 2

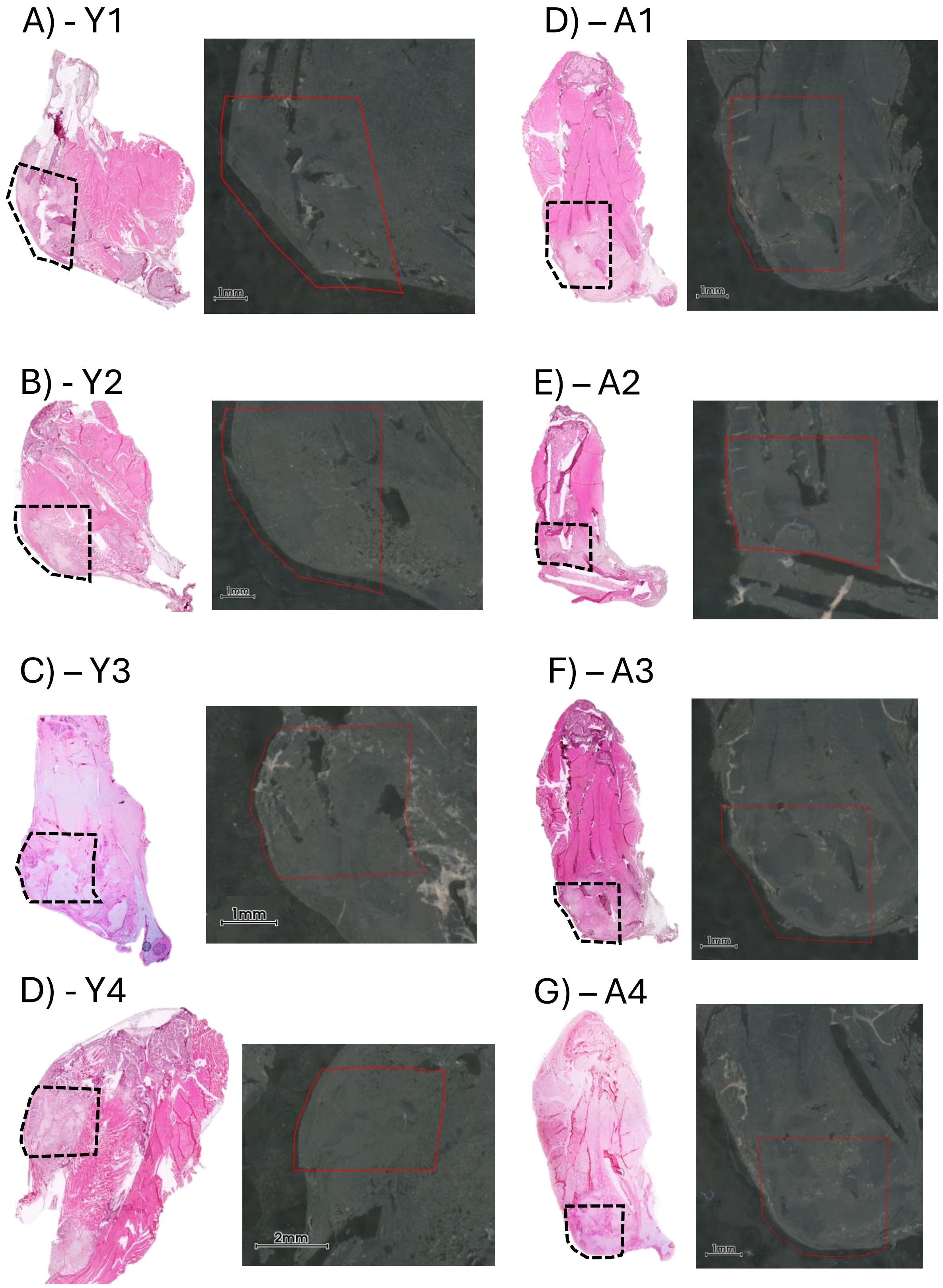


**Supplemental Figure 2: Fracture Callus Region Identification.** Hematoxylin and Eosin (pink, left) histologic staining and unstained, enzymatically digested tissue sections (right, grey) of unfixed fracture calluses from (A) young (3 mo.) and (B) aged (18 mo.) mice 10 weeks post tibial fracture showing regions of tissue selected for MALDI-MSI (black dotted box, left or red solid region, right).
