## Supplementary material for "Age-related Delays in Osteochondral Remodeling of Fracture Healing Illustrated by Mass Spectrometry Imaging": Suppl Figure 3

### Supplemental Figure 3

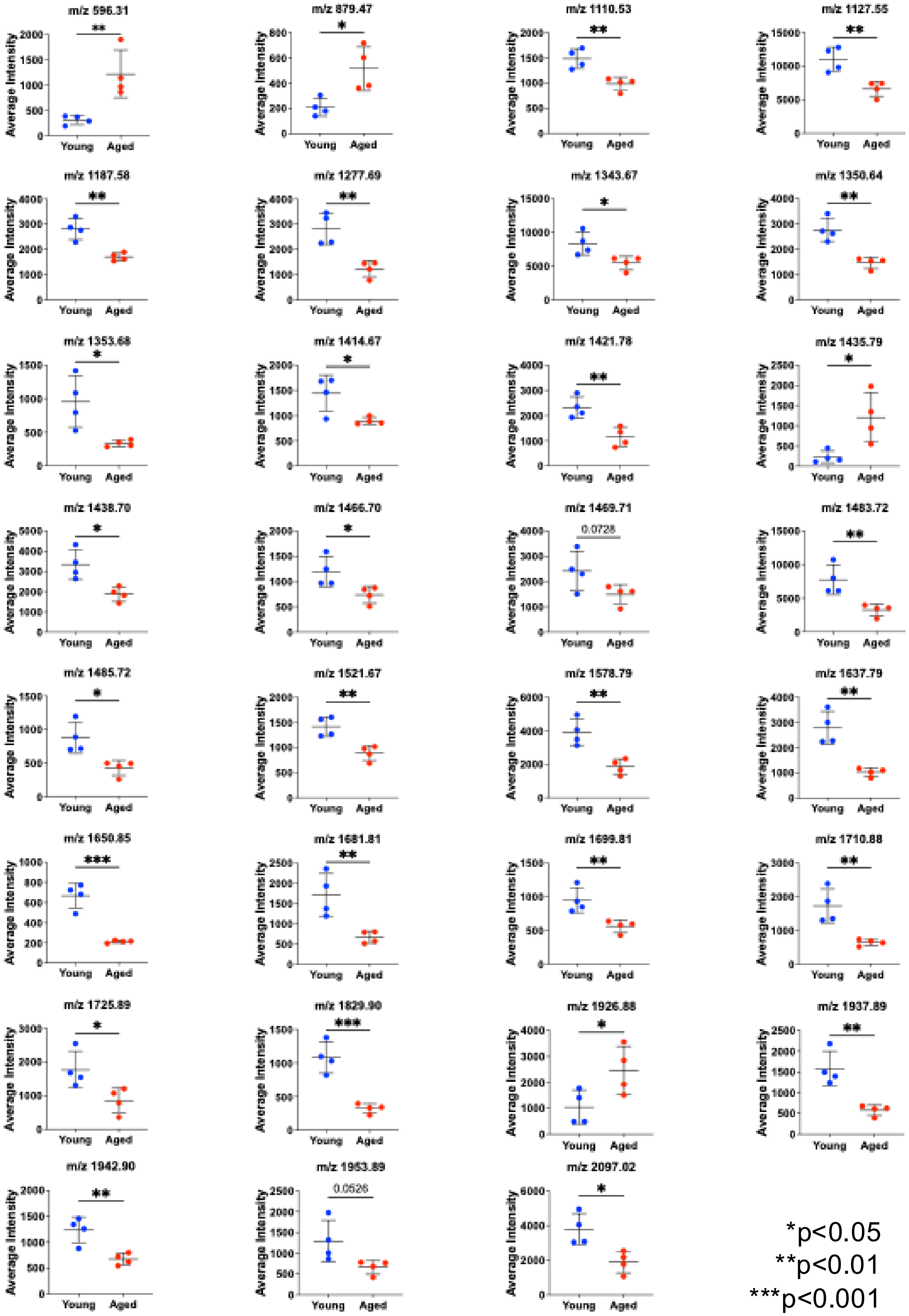

**Supplemental Figure 3: Discriminating Features Analysis by Age**: SCiLS lab identified 31 unique m/z features with an |AUROC| > 0.7 that had significant predictive ability for either the young or aged fracture callus. Exported intensities for each feature from each tissue section are compared showing cohort level significant difference in a t-test for 29 of 31 features demonstrating the consistency of regulation of each feature by age. The remaining two features trended toward significance with a p value <0.1. *p<0.05 **p<0.01 ***p<0.001
