## Supplementary material for "Age-related Delays in Osteochondral Remodeling of Fracture Healing Illustrated by Mass Spectrometry Imaging": Suppl Figure 4

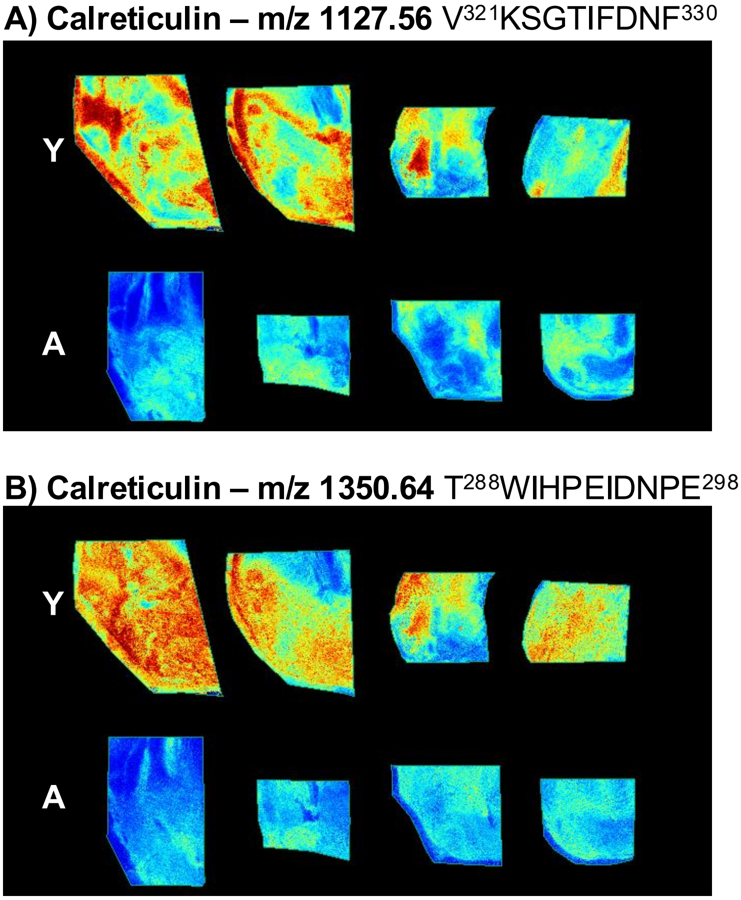


**Supplemental Figure 4: Spatial Distribution for Calreticulin as regulated by Age in Fracture Callus.** Ion intensity heatmaps for the two peptides of Calreticulin, (A) V$^{321}$KSGTIFDNF$^{330}$ m/z =1127.56 and (B) T$^{288}$WIHPEIDNPE$^{298}$ m/z = 1350.64 were identified as upregulated in the young callus group and largely shared a similar spatial distribution, further indicating each unique peptide stemmed from the same protein.
