## Supplementary material for "Age-related Delays in Osteochondral Remodeling of Fracture Healing Illustrated by Mass Spectrometry Imaging": Suppl Figure 5

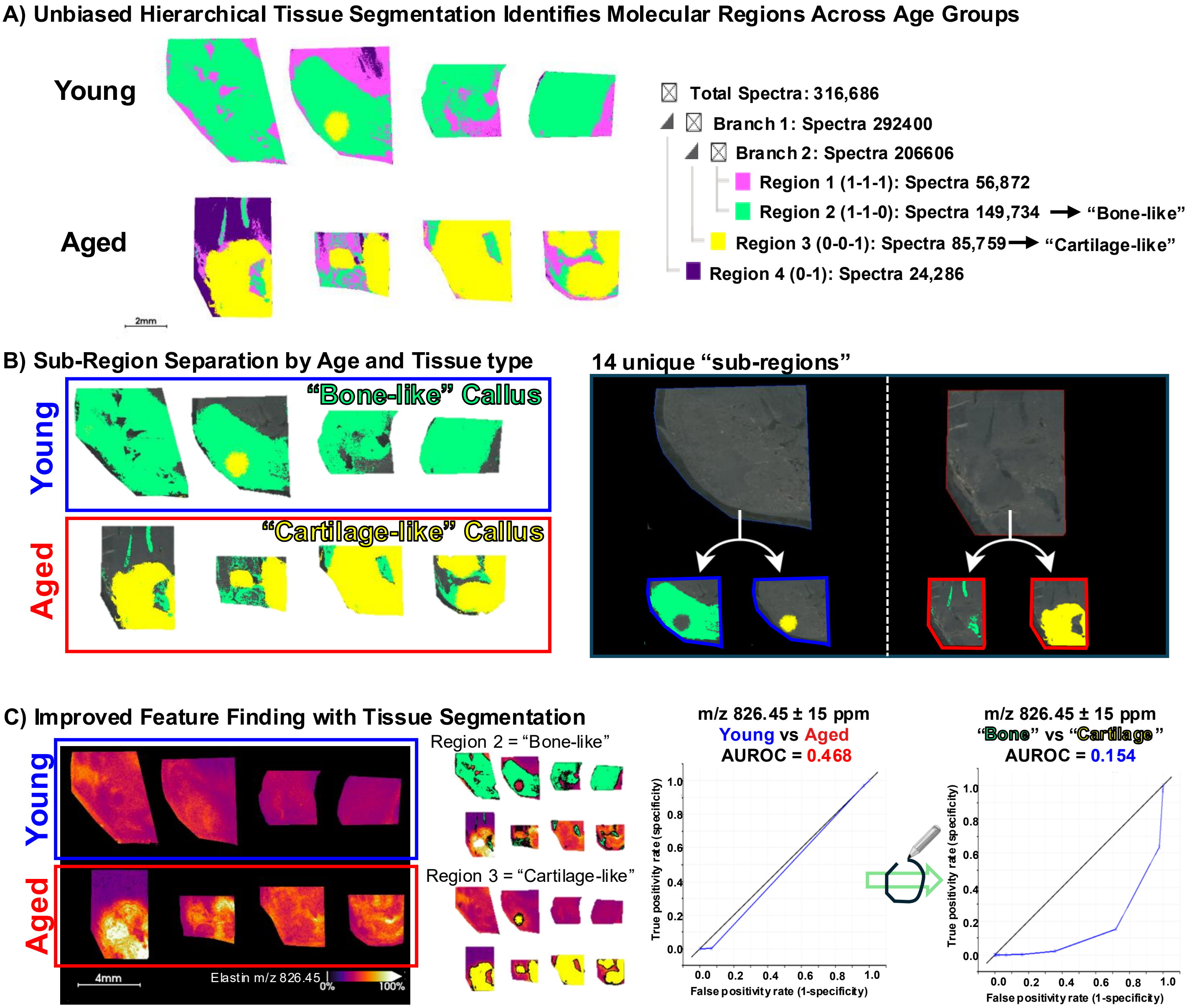


**Supplemental Figure 5: Tissue Classification by Automated Segmentation.** A) Hierarchical segmentation based on molecular signal from MALDI-MSI on fracture callus tissue samples identified several distinct tissue types. The two largest regions matched areas from H&E staining that identified more "one-like" (Region 2, green) and "cartilage-like" (Region 3, yellow) tissues areas across both young and aged callus groups. **B)** Sub-region separation using the dual labels of age-group (young, aged) and tissue type (bone or cartilage-like) resulted in 14 unique ROIs for feature extraction. **C)** Utilizing the tissue-specific ROIs, instead of age-group alone, resulted in higher AUROC predictive scores for several peptides increasing the number of identified candidates for validation, as exhibited by the identified peptide for Elastin $G^{340}GAGAIPGIGG^{350}$ at m/z 826.45 that had an AUROC value increase from a non-significant/non-predictive score by age of AUROC = 0.468 to a significant AUROC = 0.154 (|AUROC| = 1 - 0.154 = 0.846) when using the tissue segmentation.
