## Supplementary material for "Age-related Delays in Osteochondral Remodeling of Fracture Healing Illustrated by Mass Spectrometry Imaging": Suppl Figure 6

### Supplemental Figure 6

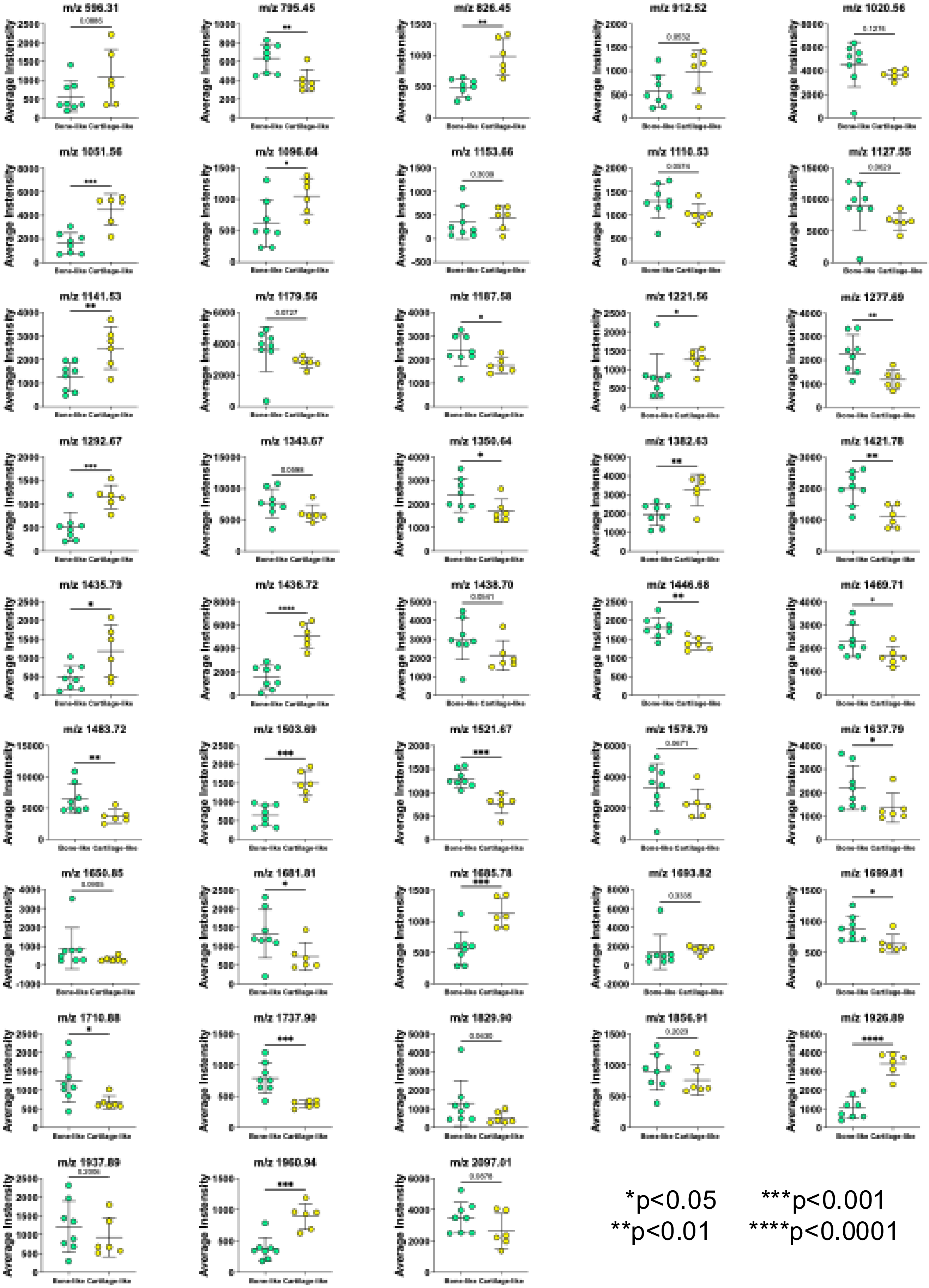

**Supplemental Figure 6: Discriminating Features Analysis by Tissue Type**: SCiLS lab identified 43 unique m/z features with an |AUROC| > 0.7 that had significant predictive ability for the automatically generated tissue regions termed “Bone-like” (Fig. 5B, green) and “Cartilage-like” (Fig. 5B, yellow). Exported intensities for each feature from each of the 14 identified tissue sub-regions are compared showing cohort level significant difference in a t-test for 27 of 43 features. Of the remaining features, 11 features trended toward significance with a p value <0.1. *p<0.05 **p<0.01 ***p<0.001
