## Supplementary material for "Age-related Delays in Osteochondral Remodeling of Fracture Healing Illustrated by Mass Spectrometry Imaging": Suppl Figure 7

### Supplemental Figure 7

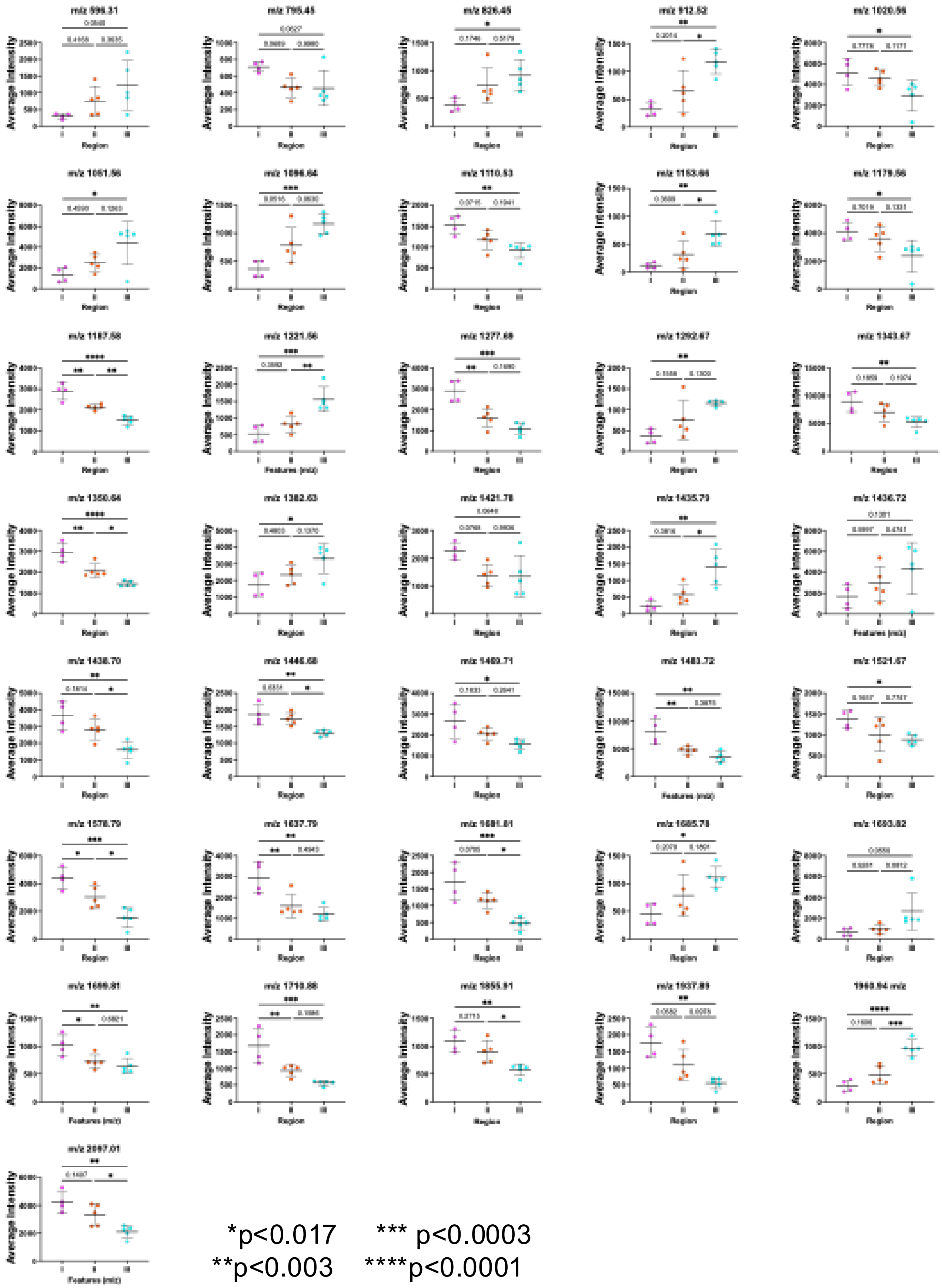

*p<0.017 *** p<0.0003
**p<0.003 ****p<0.0001

**Supplemental Figure 7: Regulation of Identified Features after Gradient Re-classification:** After hierarchical clustering sorted the 14 exported tissue sub-regions subregions into 3 new tissue classes showing a gradient of healing/matrix remodeling throughout fracture healing, 36 of the 43 features found by Discriminating Feature analysis showed a significant difference in one-way ANOVA when utilizing each new class (I, II, III) as factors. 3-way Bonferroni corrected post-hoc pairwise comparisons are shown for features that showed significant difference in initial ANOVA testing. *$p<0.017$, **$p<0.003$, ***$p<0.0003$, ****$p<0.0001$.
