## Supplementary material for "Age-related Delays in Osteochondral Remodeling of Fracture Healing Illustrated by Mass Spectrometry Imaging": Suppl Figure 8

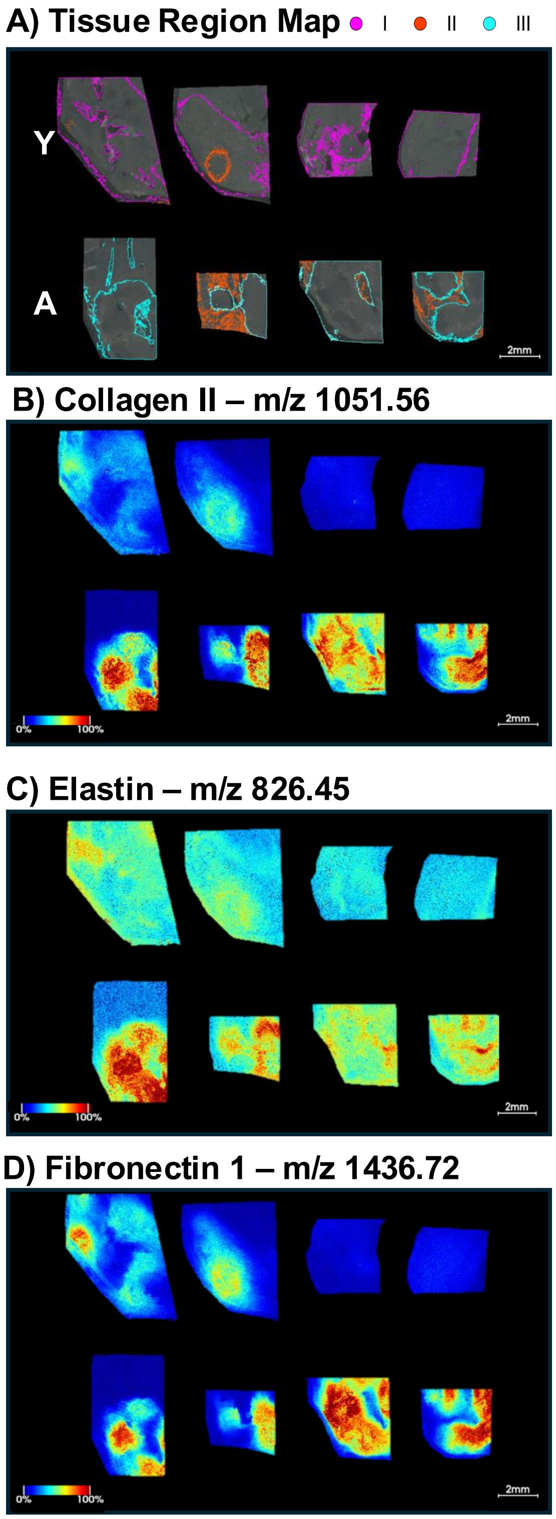


**Supplemental Figure 8: Select Ion Heatmaps for Features Identified by MALDI-MSI with Tissue Type Regulation.** Ion intensity heatmaps for three peptides identified with differential abundance across different tissue regions (A) within the fracture callus. B) Collagen II peptide $G^{1101}$PAGARGIAGP$Q^{1112}$ m/z = 1051.56, C) Elastin peptide $G^{340}$GAGAIPGIG$G^{350}$ m/z 826.45 and D) Fibronectin peptide $F^{1584}$TVPGSKSTATIN$N^{1597}$ m/z 1436.72 show higher abundance in the delayed (III) region of the callus compared to the intermediate (II) or more healed (I) regions.
